# Ankh-score produces better sequence alignments than AlphaFold3

**DOI:** 10.1101/2025.09.03.674029

**Authors:** Julia Malec, Karina Rusen, G. Brian Golding, Lucian Ilie

## Abstract

Protein sequence alignment is one of the most fundamental procedures in bioinformatics. Due to its many downstream applications, improvements to this procedure are of great importance. We consider two revolutionary concepts that emerged recently as candidates for improving the state-of-the-art alignment methods: AlphaFold and protein language models such as Ankh, ProtT5 or ESM-C. Alignment improvements can come from the structural alignment of AlphaFold-predicted structures or the scoring based on the similarity of protein embeddings produced by the protein language models. Thorough comparison on many domains from BAliBASE and CDD demonstrates that the Ankh-score method produces much better sequence alignments than the structural alignments using US-align of AlphaFold3-predicted structures. Both are better than the traditional method using BLOSUM matrices. This suggests that Ankh embeddings may possess certain information that is not available in the AlphaFold3-predicted structures. The alignment software is freely available as a web server at e-score.csd.uwo.ca and as source code at github.com/lucian-ilie/E-score.

## 1. Introduction

Protein sequences contain information necessary for the many functions they perform, which makes protein sequence alignment one of the most fundamental procedures in bioinformatics. Improvements to the quality of alignments are essential for their many downstream applications. Two concepts emerged recently as candidates for improving the state-of-the-art alignment methods. The first is AlphaFold, which, through its high quality predicted structures, can produce sequence alignments via alignment of the structures. The second is protein language models, which can help produce improved amino acid scoring schemes. Our goal is to provide a thorough comparison of these methods as well as the traditional BLOSUM matrices.

Two revolutions have been undergoing in bioinformatics and biology in recent years. The first one is AlphaFold [16], which produced a much better solution to the fundamental problem of structure prediction, accelerating drug discovery, enzyme engineering, protein-protein interaction studies, etc. AlphaFold illuminated half of the dark human proteins [3]. The second one concerns protein language models (PLMs), such as ProtT5 [7], ESM2 [21], or Ankh [6]. PLMs are trained on vast unlabelled sequence databases, such as UniProt [32] or BFD [30], to provide contextual representations for amino acid residues in the form of high-dimensional vectors, called embeddings, that capture functional and evolutionary patterns, helping to create state-of-the-art solutions for many problems in proteomics, such as structure prediction [29, 34, 16], function prediction [18, 8, 19], interaction site prediction [23, 14], etc. AlphaFold itself uses a form of embedding (Evoformer) to make its own predictions.

Because evolutionary pressure is higher on the structure than it is on the sequence, Rajapaksa et al [28] claimed that structure alignments are significantly more reliable than sequence-only alignments. Lesk and Konagurthu [20] suggest that applying AlphaFold to the sequences and aligning the predicted structures produces better sequence alignments than traditional methods based on the Needleman-Wunsch algorithm [26] with affine gap penalties [9]. They argue that “structure-based methods will for most cases supersede classical sequence-only-based methods for the central and fundamental problem of alignment.”

On the other hand, the cosine similarity of embeddings associated with amino acid residues has been used recently as a successful scoring scheme for alignment computation by several studies: pLM-BLAST [17], vcMSA [25], PEbA [15] and E-score [2, 22]. In particular, PEbA and E-score show that PLM-based scoring schemes produce much better pairwise alignments than traditional BLOSUM matrices.

Thus, we have two very promising candidates for replacing the traditional methods for computing sequence alignments. We present here a thorough comparison of the three methods: BLOSUM matrices [12], AlphaFold3 [1] (with US-align [36]), and Ankh-score [2, 22]. We use a wide variety of alignments selected from BAliBASE [31] and the Conserved Domain Database [24] and employ four different distance metrics to compare the results with the reference alignments.

The conclusion is that Ankh-score is clearly the best, followed by AlphaFold3 and then the traditional BLOSUM matrices. We provide also examples showing the superiority of the information provided by the contextual embeddings over the alignment of AlphaFold3-predicted structures.

In addition to identifying the best method for computing protein sequence alignments, we have two important observations. First, it appears that PLMs, especially Ankh, may possess certain information that is not available in the AlphaFold3-predicted structures. Second, our testing appears to indicate that the sequence alignments produced from experimentally determined structures are slightly inferior to those derived using AlphaFold-predicted structures, an intriguing hypothesis that remains to be investigated.

## 2. Material and Methods

### 2.1 Alignment methods

We compare three methods for computing protein sequence alignments. The first is dynamic programming [26] with affine gap penalties [9] using the score provided by the BLOSUM matrices [12]. The common matrices are considered: BLOSUM45, BLOSUM50, BLOSUM62, BLOSUM80, and BLO-SUM90. For most tests, BLOSUM45 gives the best results and, therefore, most of the analysis focuses on this one out of the five matrices.

The second method is the sequence alignment induced by aligning the structures predicted by AlphaFold3 [1] using US-align [36, 35]; we shall call this method *AF3US*. The idea is that the residues from the AlphaFold3-predicted structures that are aligned to be in close proximity by US-align become aligned as well in the sequence alignment. We prove US-align to be the best program for this task.

US-align, the successor of TM-align [36] reports two TM-scores, one for each protein sequence, as a measure of structural similarity. The TM-score is length independent. A score above 0.5 suggests very good alignment, the two structures sharing the same overall fold, while a score below 0.2 suggests that the proteins are likely unrelated. If two proteins of very different length are aligned, then the TM-score for the longer one will always be low, reflecting the unaligned remainder of the larger structure. In this case, it is possible that the TM-score for the shorter one is very high, meaning that the shorter protein aligns very well with a part of the longer one.

The third method employs also dynamic programming with affine penalties but uses for scoring the similarity of Ankh embeddings [6]. Precisely, the Ankh-score [2, 22] for two residues *a*_1_ and *a*_2_ is given by the cosine similarity between the vectors *v*_*i*_ = *Ankh*(*a*_*i*_), *i* = 1, 2, computed by Ankh for the two residues, that is:

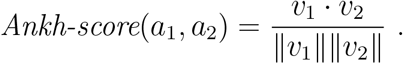

We prove Ankh to be the best PLM for this task.

### 2.2. Testing datasets

We have selected a wide variety of domains from BAliBASE [31] and the Conserved Domain Database (CDD) [24]. We attempted to cover many different identity levels, as a main criterion that influences the quality of the alignments. Figure 1 gives the list of 20 domains we use from each database, with the identity distribution and the number of sequences. More details are provided in the Supplementary Tables 1-2; see also the ‘Data’ sheet of the Supplementary excel file.

**Figure 1:**
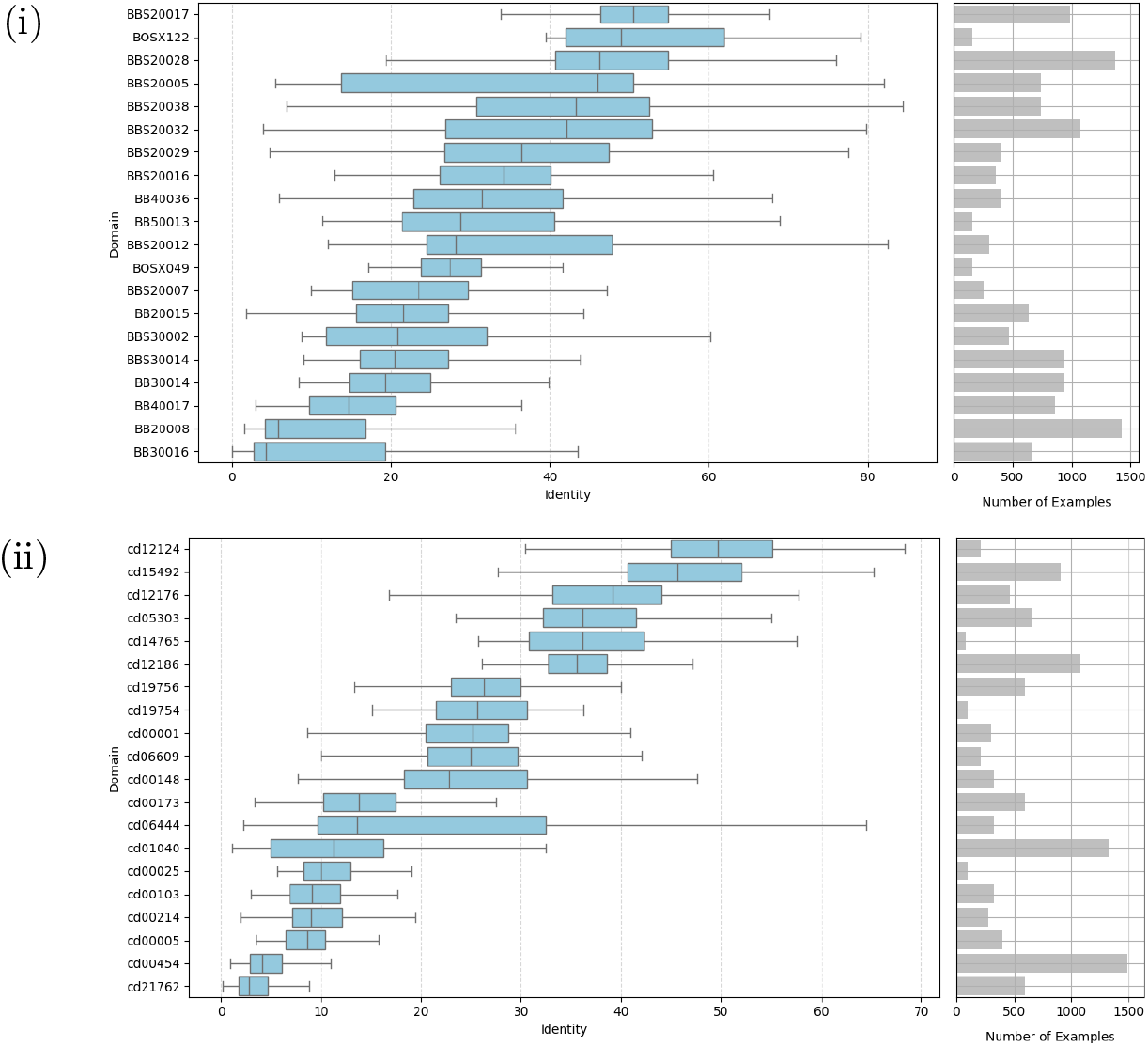
Testing domains from (i) BAliBASE and (ii) CDD. For each domain we plot the identity distribution as box-and-whiskers on the left and the number of sequences on the right. Further details are given in the Supplementary Tables 1-2.

### 2.3. Comparison procedure

Each domain consists of a number of protein sequences, say *n*, and a reference multiple sequence alignment (MSA) of all *n* sequences. A test is any pair of sequences, 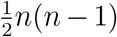 in total for a domain with *n* sequences. The two sequences have a reference alignment induced by the original MSA. This is the reference alignment with which all computed alignments between the two sequences are compared.

For each method, an alignment is computed and its distance to the reference alignment is measured according to one of several distance metrics. We have used four different distances. The first one is the inter-alignment distance [28], d_ia_, and measures the area between the alignments paths in the dynamic programming matrix. The second one is the relative displacement distance [2], d_d_, and consists of all position differences of all residue pairs. The third one, d_cc_, measures the distance to the closest position with the same context [2]. The last one, d_pos_, comes from three distances, d_ssp_, d_seq_, d_pos_, which are true metrics inspired from the sum-of-pairs score, differing from each other in the way they consider gaps [4]. As d_pos_ considers the most information about gaps – the sequence and the position of the gap in the sequence – it is the most relevant of the three. In order to reduce the complexity of the comparison, we use only d_pos_ out of the three.

For each method tested on a domain and evaluated according to a distance, we report (i) the average distance from the alignments computed by the method to the reference alignments and (ii) the number of tests won by the method. The methods are compared in pairs, enabling the use of Wilcoxon signed-rank test; we consider p-values smaller than 0.01 significant. When calculating the number pf tests won by a method we consider only statistically significant tests. In the Supplementary excel file that contains the comparisons, P-values larger than 0.01 are shown in red.

### 2.4. PLM selection

We considered several top PLM candidates for producing the best sequence alignments: ProtT5; ProstT5 [11]–a powerful fine-tuning of ProtT5 using structures; ESM-C–the successor to ESM2; and Ankh. In order to show that Ankh is the best method, we compared it with the other three. A summary of the results is shown in Figure 2. Recall that only statistically significant tests are considered, therefore the total number of tests is different among the comparisons. Full results are presented in the Supplementary excel file, sheets: ‘Ankh vs ProtT5,’ ‘Ankh vs ProstT5,’ and ‘Ankh vs ESM-C.’

**Figure 2:**
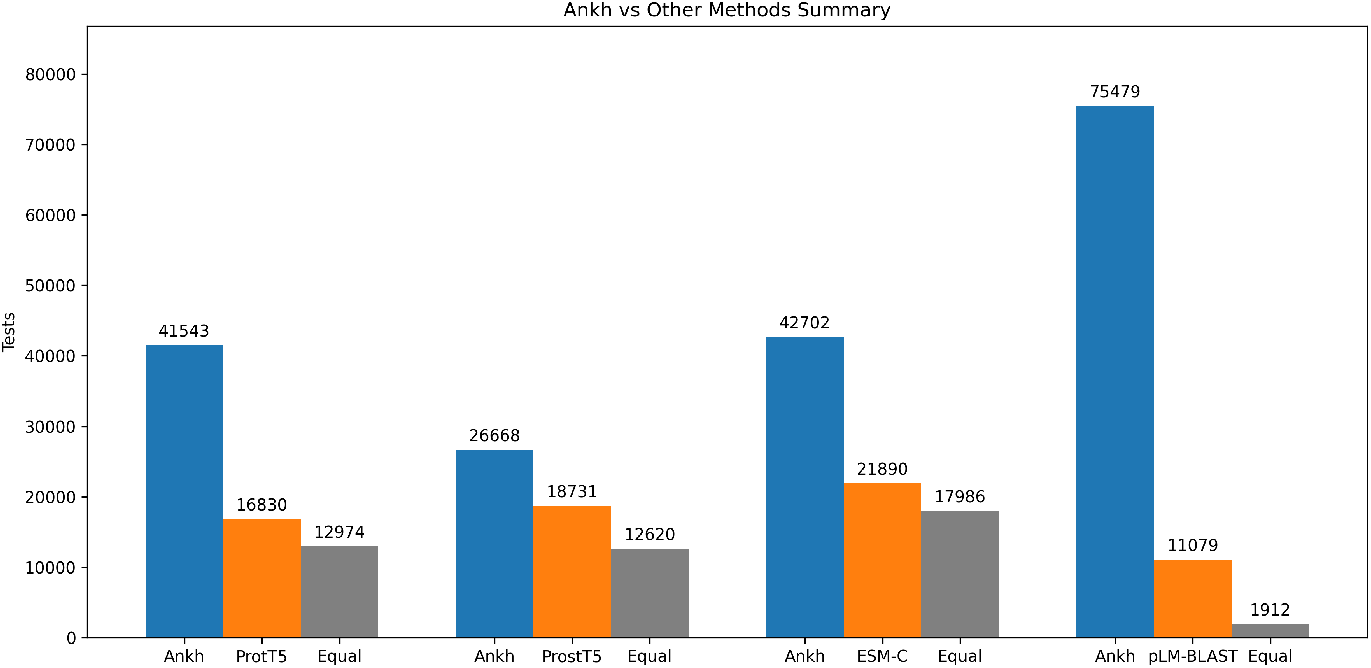
Summary of comparisons between Ankh and ProtT5, ProstT5, ESM-C, pLM-BLAST. The total number of tests won by each method is given, where only statistically significant tests are considered.

We compared as well the alignments produced using the Ankh-score with other top methods that use PLMs. One of them is PEbA which uses also dynamic programming with scores provided by the cosine similarity of ProtT5 embeddings. Therefore, our comparison between Ankh and ProtT5 indicates our method superior to PEbA. The last method to consider is pLM-BLAST and the summary of the comparison is shown in Figure 2 with full details in the sheet ‘Ankh vs pLM-BLAST’ of the Supplementary excel file. Ankh clearly outperforms all the other methods considered.

### 2.5. Gap penalty sensitivity analysis

The affine penalties we have used in our tests are -1.5 for gap opening and -0.15 for gap extension and have been determined experimentally. In order to show the robustness of our results with respect to the choice of gap penalties, we performed several tests to assess the behaviour of Ankh’s performance for penalties in the interval [ −1.7, −1.3] × [ −0.17, −0.13]. The results for the d_ia_ distance are shown as heat maps in Figure 3. It can be seen that significant variations in gap penalties have little impact on the alignment quality. The results for the other three distances are given in the Supplementary Figure 1. Similar patterns with very small variations are seen as well for these distances, indicating that the alignment computation is robust with respect to the choice of gap penalties.

**Figure 3:**
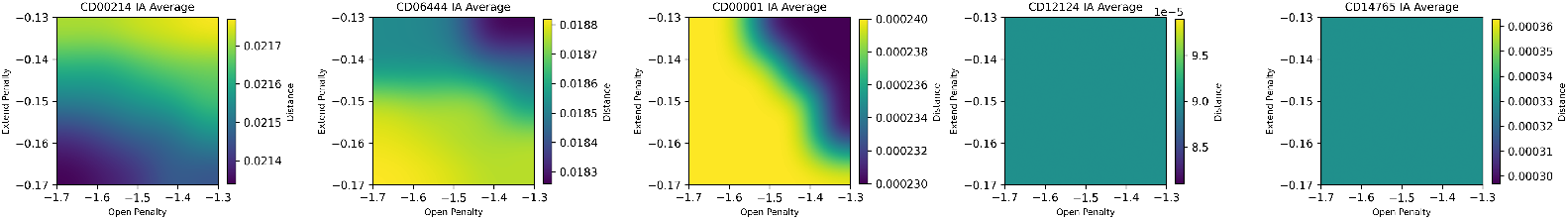
Heat maps showing the variation of the d_ia_ distance between the Ankh-score alignments and the reference for several domains.

### 2.6. Structure aligner selection

We considered three candidates for aligning structures with the purpose of extracting sequence alignments: US-align, DALI [13] and Foldseek [33]. A summary of the results is shown in Figure 4. Recall that only statistically significant tests are considered, therefore the total number of tests is different among the comparisons. Full results are presented in the Supplementary excel file, sheets: ‘US-align vs DALI,’ and ‘US-align vs Foldseek.’ US-align outperforms both methods by a significant margin.

**Figure 4:**
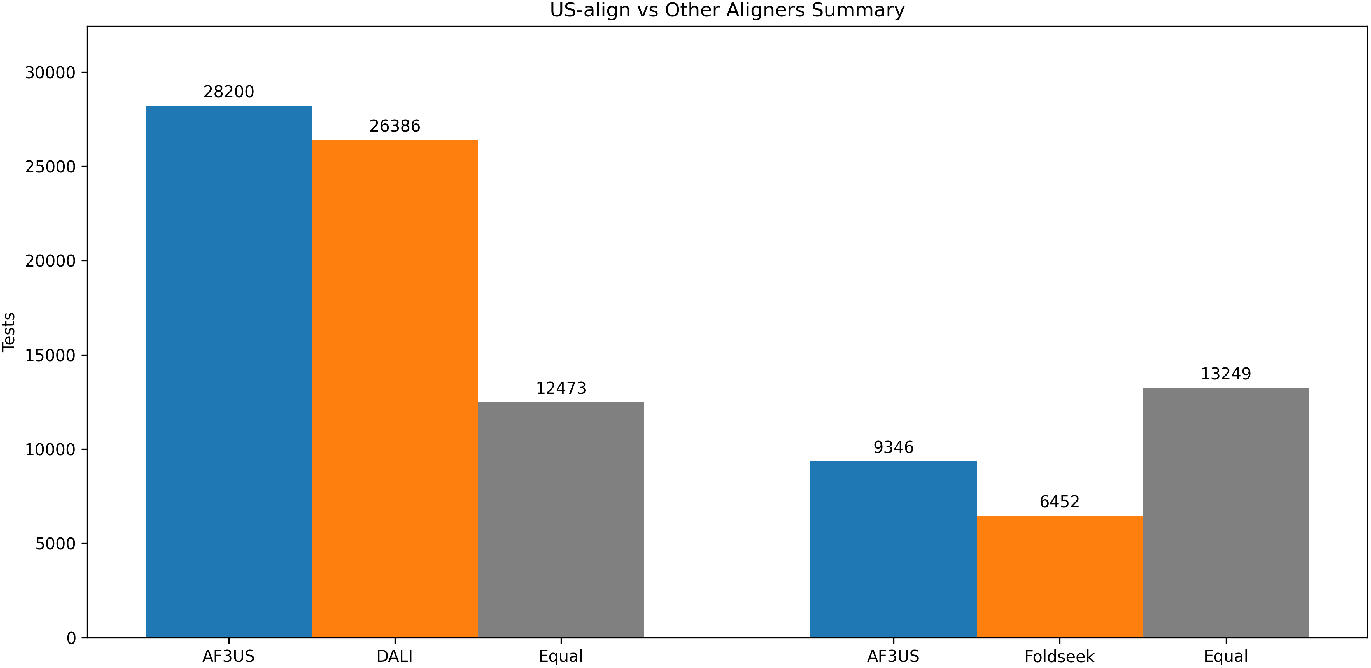
Summary of comparisons between US-align and DALI, Foldseek. The total number of tests won by each method is given, where only statistically significant tests are considered.

## 3. Results

### 3.1. All-against-all comparison

We first compare all methods together, separately on BAliBASE and CDD datasets. The comparison is shown in Figure 5 for the d_ia_ distance.

**Figure 5:**
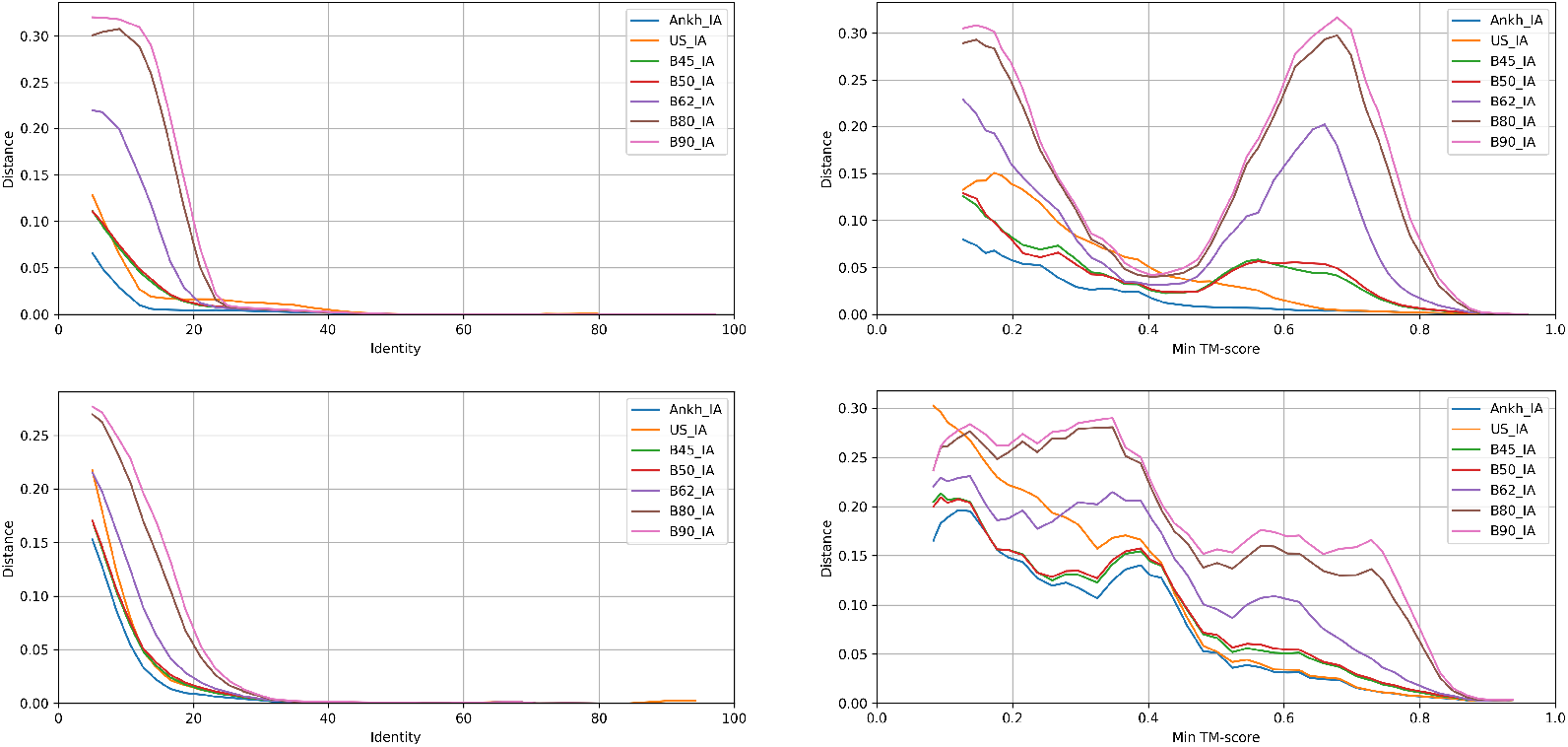
All methods are compared on all tests for the d_ia_ distance. The plots in the first row are for BAliBASE datasets, while those is the second row are for CDD. In the first column, the scores are sorted by the identity level of the sequences, while in the second column, they are sorted by the minimum TM-score.

Similar trends are exhibited by the other distances, as shown in Supplementary Figures 2-4. All methods are plotted: BLOSUM45, BLOSUM50, BLOSUM62, BLOSUM80, BLOSUM90, AF3US, and Ankh-score. The top row corresponds to the BAliBASE datasets and the bottom one to the CDD datasets. In each row, in the first plot the tests are sorted by identity level, whereas in the second one they are sorted by minimum TM-score. Identity level and minimum TM-score are the most important parameters, as sequences with higher identity level are easier to align, and higher TM-score indicates better performance for the structural alignment.

For each plot, in order to have good control over which data is being used to draw the line, the average distance of all tests in a sliding window of fixed size is computed and the window is shifted by a fixed step. For the identity plots, the window size is 10 and the step is 2. For the min TM-score plots, the window size is 0.1 and the step is 0.02.

The first observation, common to all plots, is that the Ankh-score alignments are always the best. The BLOSUM matrices with lower index perform better, BLOSUM45 being the best with BLOSUM50 a very close second. For that reason, from now on, when discussing the behaviour of BLOSUM matrices, we will restrict our attention exclusively to BLOSUM45.

AF3US has an interesting behaviour. It starts the worst (compared to Ankh-score and BLOSUM45) and then it gradually becomes second best, behind Ankh-score. This happens in the identity plots around identity level 10-15% and in the min TM-score plots around TM-score 0.5. After that, in the min TM-score plots it remains second best until the end, inching closer and closer to the performance of the Ankh-score algorithm. In the identity plots, the behaviour is somewhat different. The distances for all methods eventually become very close to zero, except for the AF3US, which is sometimes hovering above the abscissa, even for very high identity values. This is more pronounced in the BAliBASE plots, but visible also in the CDD plots. This behaviour may be partly explained by the tests with TM-score below 0.5, which may cause poor alignments, thus increasing the average distance. However, as we discuss below, this is not the only factor.

As structural alignments are considered reliable when the TM-score is above 0.5, we have plotted separately the results including only tests with minimum TM-score above 0.5 in Figure 6; only the d_ia_ distance is shown but similar trends are exhibited by the other distances, as shown in Supplementary Figures 5-7. The performance of AF3US is improved compared with the case of all tests. It now starts as second best, behind Ankh-score, from the beginning. However, the “wavering” behaviour is still present. We have provided zoom inserts to the identity plots in Figure 6 to make this more clearly visible. In the BAliBASE plots, AF3US is the only one displaying this behaviour, whereas in the CDD plots all methods appear to falter to some degree. Finally, we notice that Ankh-score alignments are still the best, even when the tests with minimum TM-score below 0.5 are removed, which greatly favours AF3US.

**Figure 6:**
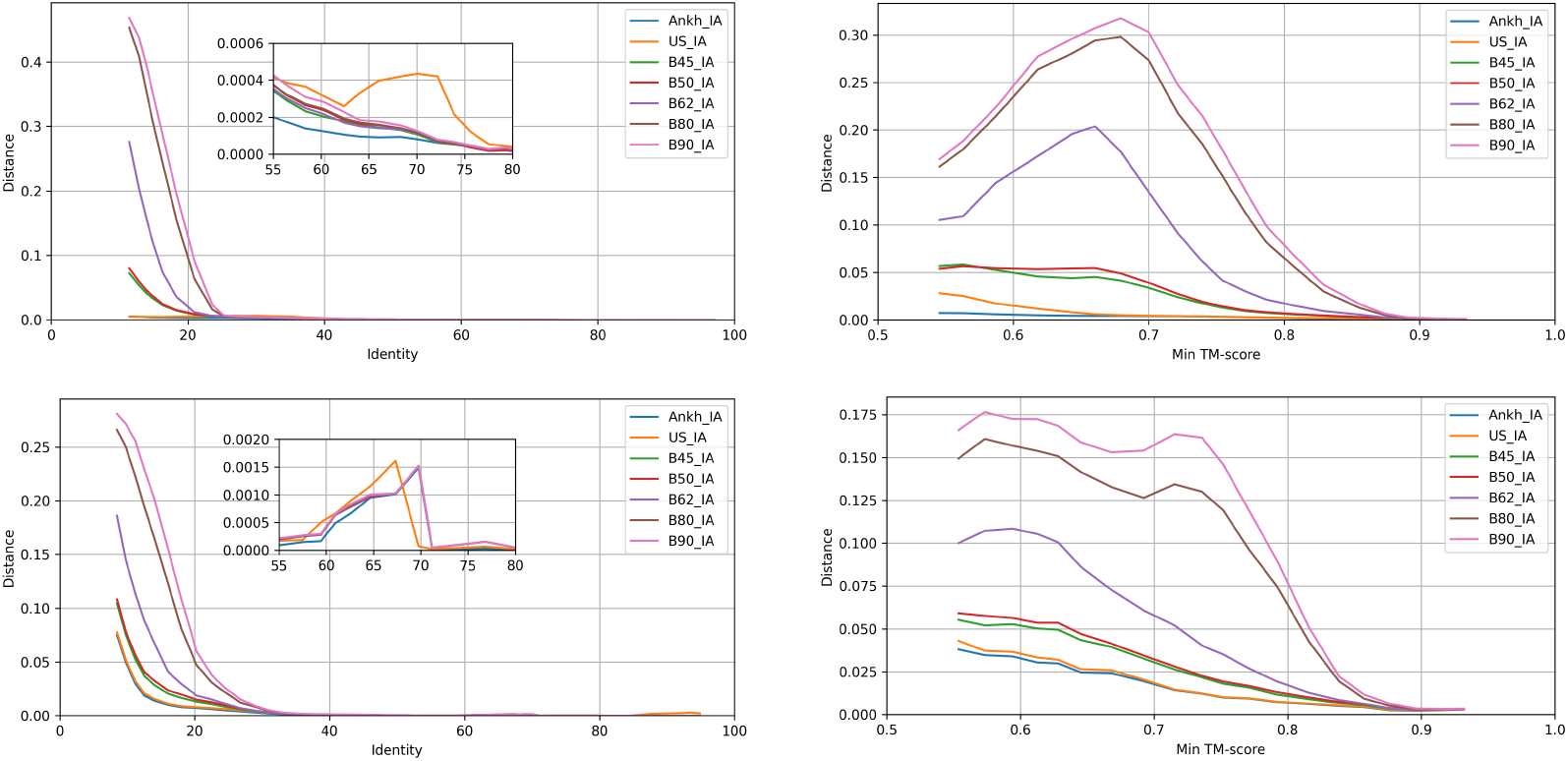
All methods are compared on tests with minimum TM-score *>* 0.5 for the d_ia_ distance. The plots in the first row are for BAliBASE datasets, while those is the second row are for CDD. In the first column, the scores are sorted by the identity level of the sequences, while in the second column, they are sorted by the minimum TM-score.

### 3.2. Head-to-head comparison

A more precise comparison is done by considering two methods at the time and computing the statistical significance, using Wilcoxon signed-rank test, of the comparison on each individual domain. We compare Ankh-score, AF3US and BLOSUM45 in pairs. Figure 7 gives a visual presentation for all comparisons in the form of dot plots. The Ankh-score dominates AF3US clearly, and even more so BLOSUM45. The performance of AF3US and BLOSUM45 appear very close in the all-tests plots, but AF3US dominates BLOSUM45 in the high TM-score plots.

**Figure 7:**
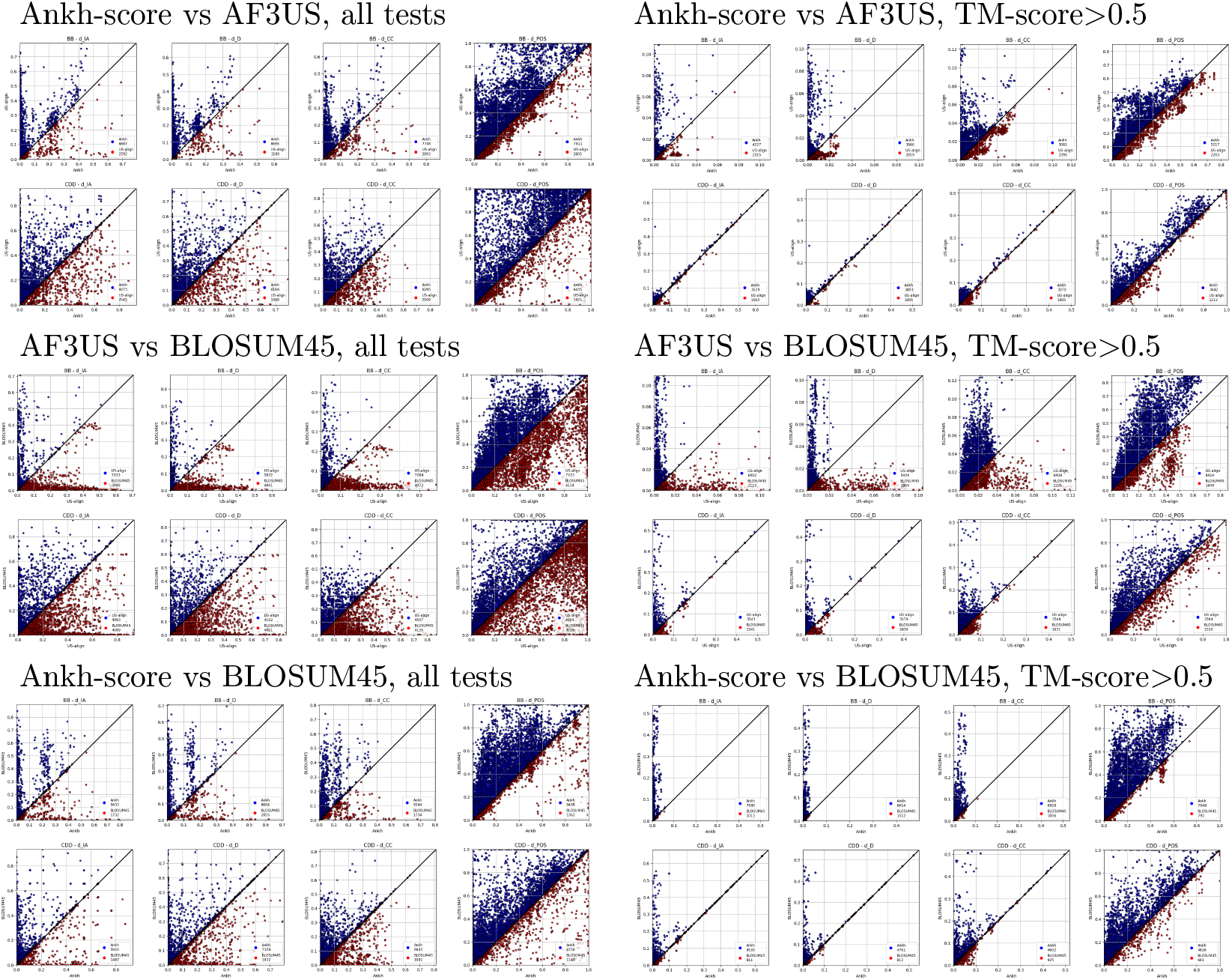
Dot plots comparing the three methods pairwise, as indicated in the table. In each group the first row is for BAliBASE and the second for CDD database, while the four columns correspond to the distances d_ia_, d_d_, d_cc_, d_pos_, resp.

A summary of results when all tests are considered, irrespective of the TM-scores, is given in Supplementary Table 3. The table gives separate results for each of the four distances considered: d_ia_, d_d_, d_cc_, d_pos_, resp. It has three sections, one for each pair of methods considered. In each case, the number of domains won and number and percentage of tests won is given, separately for each database, BAliBASE and CDD, that is, by summing/averaging the individual results for the domains from each database, as well as combined. When the number of domains won by each method is given, “null” represents the number of domains for which the result is not statistically significant. When the number and percentage of tests won is given, “equal” represents the fraction of tests for which the performance of the two methods was identical.

While different distances present slightly different results, they are consistent, revealing similar trends. Also, BAliBASE and CDD domains behave differently, but present similar pictures. Ankh-score is always winning with a very wide margin. The winning margin is larger over BLOSUM45 than over AF3US. AF3US wins always over BLOSUM45 – with a single exception provided by the case of CDD domains, percentage of tests won for the d_d_ distance – but the winning margin is not very large. The competition is closer for the CDD domains than for BAliBASE ones.

An identically organized table but using only the tests with minimum TM-score above 0.5 is shown in Supplementary Table 4. The general trends are preserved, with AF3US performing slightly better than before, given the advantage of high TM-scores.

An overall summary of results is presented in Figure 8. The number of “null” tests is larger in the case of tests with TM-score over 0.5 due to the elimination of many tests causes the p-values to increase. The general trends mentioned above are clearly visible. Ankh-score wins 78.75% of domains against AF3US, whereas AF3US wins only 10.63%. AF3US wins 59.38% of domains against BLOSUM45, with BLOSUM45 winning 30.00%. In the case of tests with minimum TM-score above 0.5, the performance of AF3US is slightly improved but the trends are preserved.

**Figure 8:**
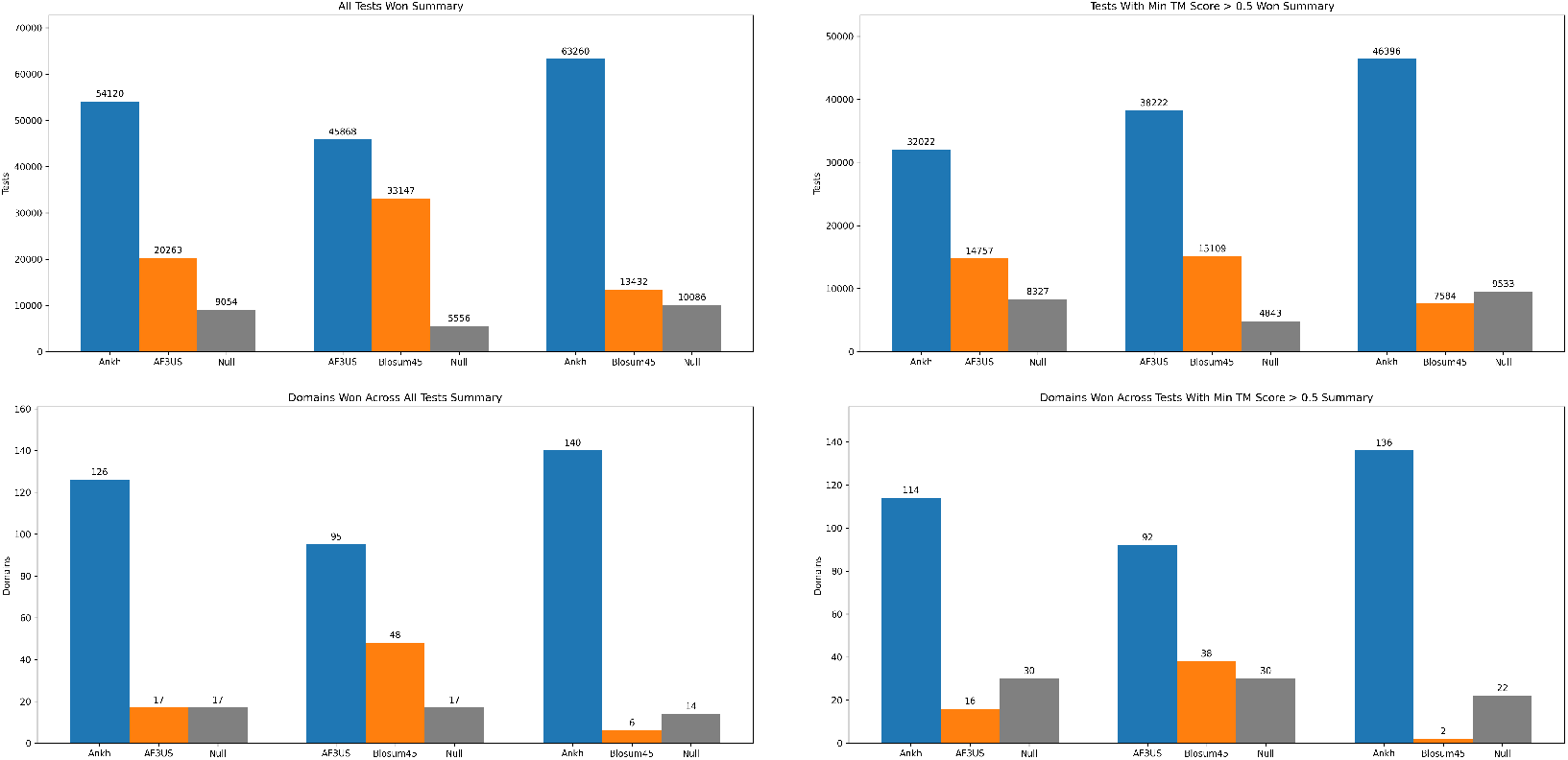
Summary of results. Total numbers of tests and domains won, combining the results for all distances and the two databases (Supplementary Tables 3-4).

Full details are given for interested readers in the Supplementary Excel file, sheets ‘Ankh vs AF3US,’, ‘AF3US vs BLOSUM45,’ and ‘Ankh vs BLOSUM45.’ Wilcoxon test p-values below 0.01 are considered statistically significant and only these tests are considered when summary results are calculated. Those with p-value above 0.01 are ignored; they are shown in red in the tables.

### 3.3. Other PLMs

We proved that Ankh provides clearly superior sequence alignments compared to those produced by the alignment of predicted structures. It is natural to ask whether only Ankh has this property or other PLMs do as well. We have compared the other PLMs against AF3US and, even though not equally, they all surpass AF3US. The summary results are shown in Figure 9 with full details given in the Supplementary Excel file, sheets ‘ProtT5 vs AF3US,’ ‘ProstT5 vs AF3US,’ and ‘ESM-C vs AF3US.’ As expected, Ankh wins by the largest margin. Then we have, in decreasing order of performance, ProstT5, ProtT5, and ESM-C. ESM-C has the weakest performance but it still outperforms AF3US.

**Figure 9:**
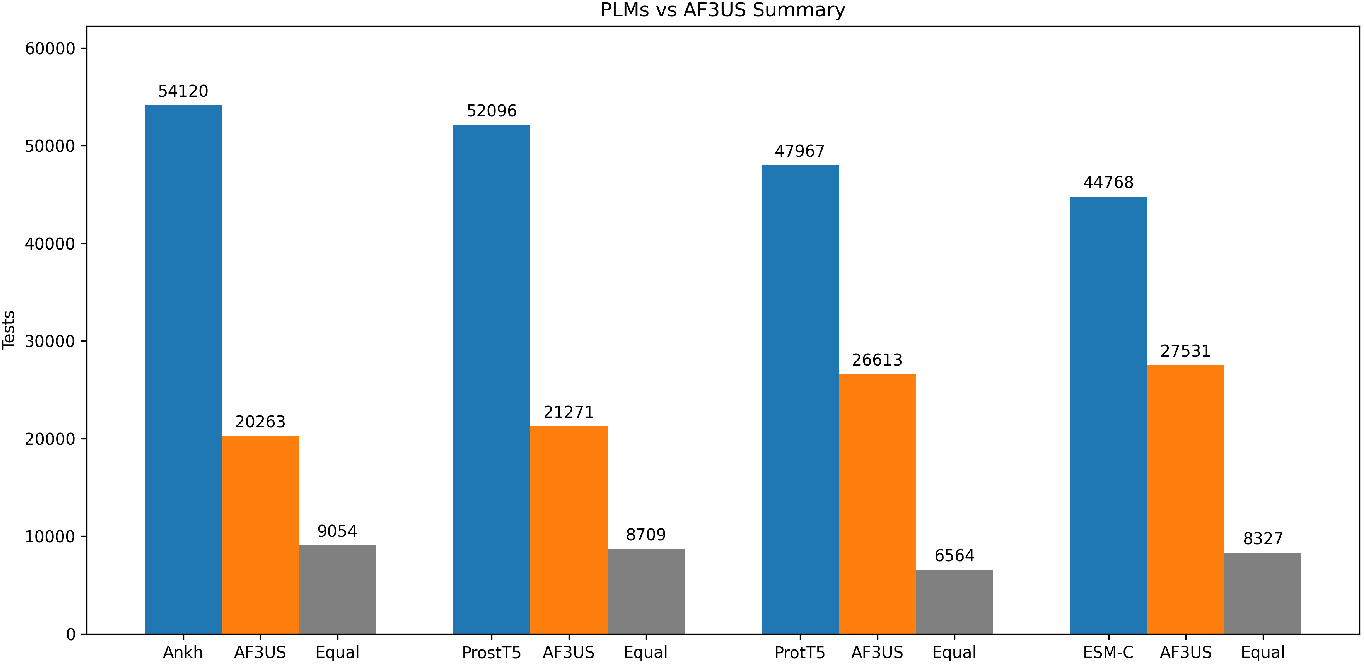
Comparison of four PLMs against AF3US. The bars indicate the number of tests won by each PLM and AF3US, resp. Only statistically significant test are considered, which explains the different number of tests in each case.

We note also that the difference between the advantages of various PLMs over AF3US is smaller than the difference between their head-to-head comparison. For example, there is a large advantage of Ankh over ProstT5 in the head-to-head comparison, as seen in Figure 2. However, when comparing each with Af3US, the advantage of Ankh over ProstT5 is smaller. This is true to some extent for all PLMs, as seen in Figure 9. Even more interesting is the domain comparison. Ankh wins over ProstT5 62 to 39 (see the ‘Ankh vs ProstT5’ sheet), whereas Ankh vs AF3US is 126 to 17, a slightly smaller difference than ProstT5 vs AF3US, which is 131 to 12. The explanation is that, sufficiently many times, when Ankh beats ProstT5, then they both beat AF3US, but when Ankh loses to ProstT5, it may lose also to AF3US. This happens for domains with very high TM-score such as cd00148, cd0644, BBS20028, BBS20029 (see their rows in the excel file). ProstT5 is a fine tuning of ProtT5 using structural information, therefore its strong performance on these high structural similarity domains is expected.

It may be that ProstT5 complements the information contained in Ankh embeddings, therefore some combination of the two, or fine tuning Ankh using structural information, may produce even better results.

The important conclusion of these comparisons is that they appear to indicate that PLMs in general may be able to capture certain information that is not available in the AlphaFold3-predicted structures.

## 4. Case studies

We present in this section three examples of proteins pairs with their three alignments: the reference, Ankh-score and AF3US. The proteins’ organism and length are given in Table 1. Their domains have been identified using SUPFAM (supfam.org) [10, 27].

**Table 1:** Three case studies. The protein pairs studied are given their ID, name, organism and length.

| Protein ID | Protein name | Organism | Length |
| --- | --- | --- | --- |
| 2D1K_C | WH2 domain of human Metastasis Suppressor Protein 1 (MTSS1) | *Homo sapiens* | 32 |
| 3UE5_B | Domain D of Spire protein | *Drosophila melanogaster* | 66 |
| HT16_HYDAT | Tyrosine-protein kinase HTK16 | *Hydra vulgaris* | 757 |
| 1d4t_A | SH2 domain of human SAP protein 1A | *Homo sapiens* | 104 |
| YXJL_BACSU | Uncharacterized transcriptional regulatory protein YxjL | *Bacillus subtilis* (strain 168) | 218 |
| DEGU_BACSU | Transcriptional regulatory protein DegU | *Bacillus subtilis* (strain 168) | 229 |

### 4.1. MTSS1 vs Spire

The MTSS1 protein has one domain: WH2 superfamily, Wiskott Aldrich syndrome homology region 2 (WH2 motif) found in Metastasis suppressor protein [1..31]. The Spire protein has two domains: WH2 superfamily, Wiskott-Aldrich Syndrome Homology (WASP) region 2 (WH2 motif) [5..30] and WH2 superfamily, fourth tandem Wiskott-Aldrich Syndrome Homology (WASP) region 2 (WH2 motif) [37..63]. The TM-scores are 0.51 and 0.31, which means one is borderline good, the other not very good. Yet, this is a simple test, as the protein sequences are very short. The alignments are shown in Figure 10. The reference aligns the WH2 domain in MTSS1 with the second WH2 of the Spire protein. Ankh-score does the same, producing an identical alignment with the reference. AF3US aligns the WH2 domain in MTSS1 with the first WH2 of the Spire protein, producing a completely different alignment.

**Figure 10:**
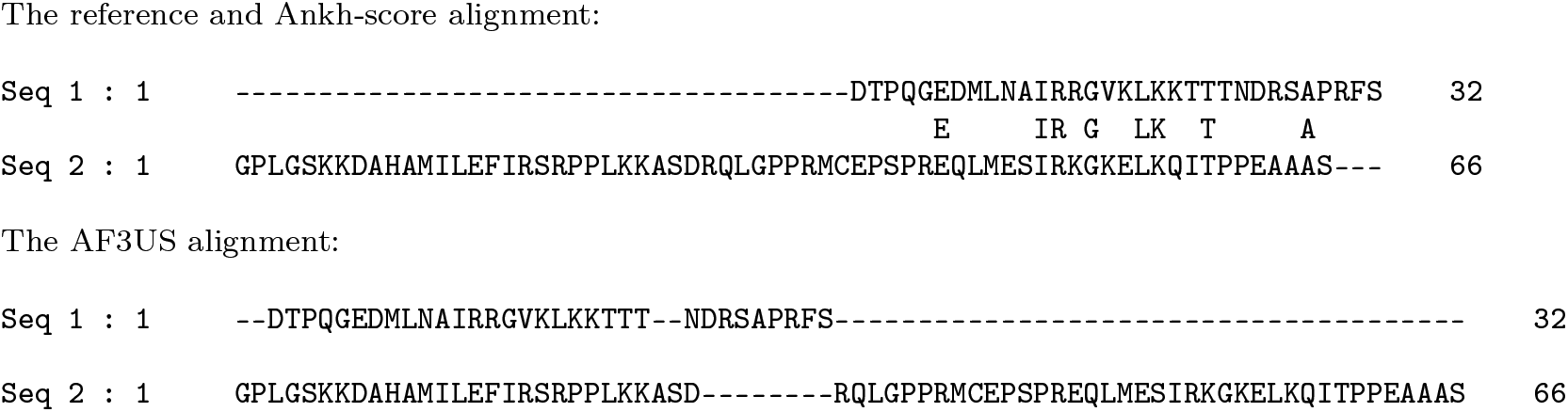
The sequence alignments for MTSS1 (`Seq 1`) and Spire (`Seq 2`) proteins. (Only two alignments are shown because the Ankh-score alignment is identical with the reference.)

For more clarity, we represent the alignments as structural alignments in Figure 11 (first row). Here we used PyMol [5] to construct structural alignments from the sequence alignments, based on the AlphaFold-predicted structures. We used the ‘pair-fit’ command of PyMol which forces a residue-by-residue alignment between the two AlphaFold-predicted structure’s backbone alpha-carbon atoms according to the sequence alignment, then finds the best rigid-body transformation, that minimizes RMSD (root mean square deviation). The result is a structural alignment that can be used to better visualize the differences between the sequence alignments. It is very clearly visible in the first row of Figure 11 that the Ankh-score alignment is identical with the reference, whereas the one by AF3US is completely different.

**Figure 11:**
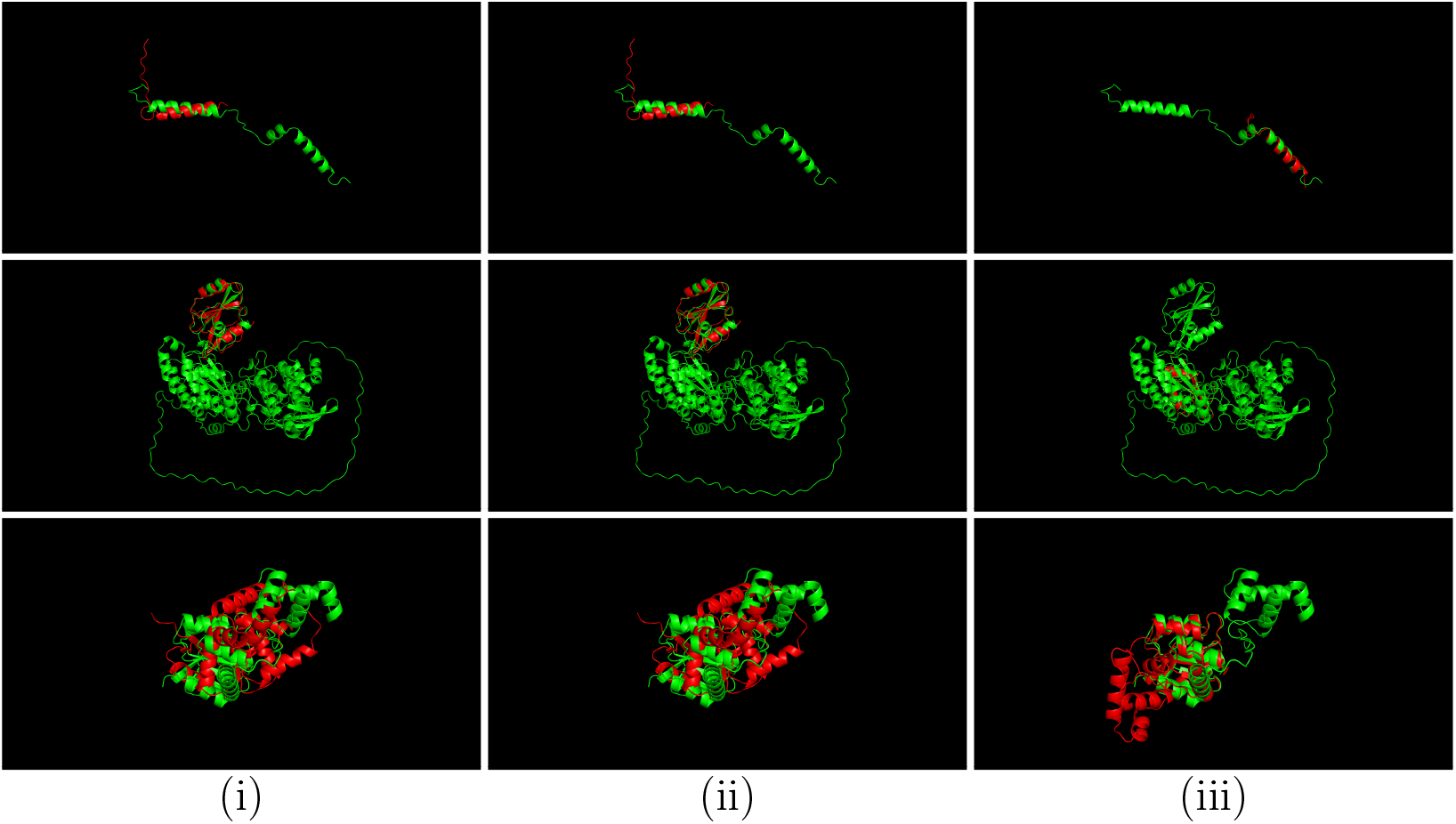
Visualizations of the three alignments: (i) reference, (ii) Ankh score, (iii) AF3US. Row 1: MTSS1 (red) and Spire (green) proteins; row 2: HT16 (green) and SH2 SAP (red) proteins; row 3: YxjL (red) and DegU (green) proteins.

### 4.2. HT16 vs SH2 SAP

The HT16 protein has four domains: SH2 (Src Homology 2 domain) [5..105], Ankryin repeat domain [99..274], another SH2 [284..503], and protein kinase, catalytic subunit [477..741]. The SH2 SAP domain has one SH2 domain [7..102]. The TM-scores are 0.84 and 0.13. These are two proteins of very different length, which means one of the TM-scores has to be low. The higher score is very high, which means good alignment of the shorter protein with a subregion of the longer. The two SH2 domains have similar shape but potentially different functions based on possible subtle differences in their surface loops, which confuses the structural alignment.

The alignments are shown in Figure 12. The SH2 domains differ in length – 101 and 220 for the first protein and 96 for the second – and the ones of similar length are aligned by the reference, that is, it aligns the SH2 domain of the second protein with the first SH2 domain of the first protein. The Ankh-score alignment does the same, being nearly identical with the reference. The AF3US alignment, on the other hand, aligns the SH2 domain of the second protein with the second, much longer, SH2 domain of the longer protein. The alignments are also visualized in second row of Figure 11.

**Figure 12:**
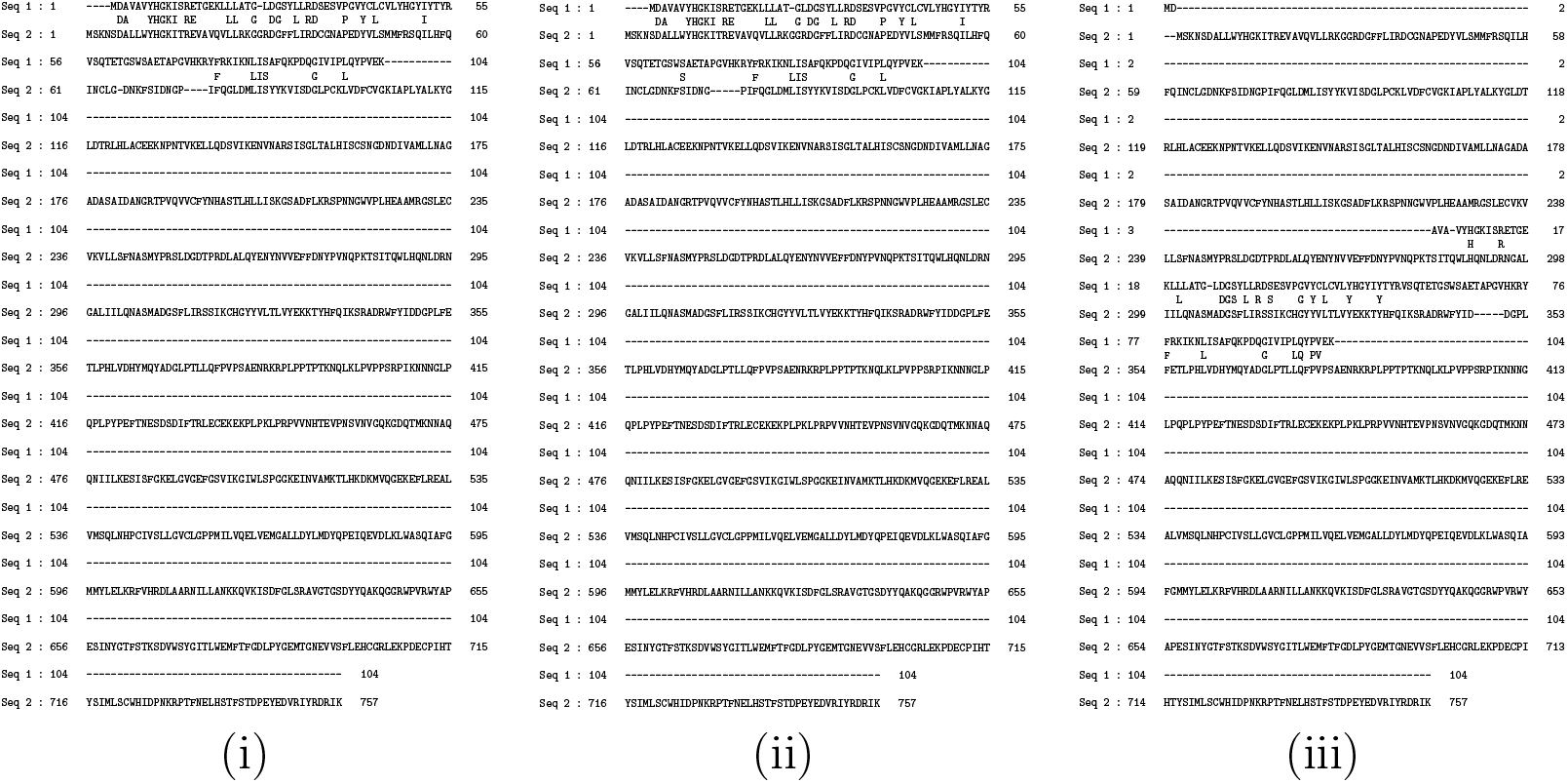
The sequence alignments for HT16 (`Seq 1`) and SH2 SAP (`Seq 2`): (i) reference, (ii) Ankh score, (iii) AF3US.

### 4.3. YxjL vs DegU

The Yxjl protein has two domains: CheY related [5..132] and GerE-like (LuxR/UhpA family of transcriptional regulators) [158..304]. The DegU protein has two domains: CheY related [3..136] and GerE-like (LuxR/UhpA family of transcriptional regulators) [149..217]. The TM-scores are 0.62 and 0.59. These are two proteins of similar lengths for which both TM-scores are over 0.5, indicating good structural alignment. They have two common domains, in the same order.

The alignments are shown in Figure 13. The reference alignment indicates, as expected, that the common domains in the two protein sequences are aligned against each other. The Ankh-score alignment is, again, identical with the reference. AF3US aligns the first domain perfectly, but then completely misses the second one. It aligns each occurrence of the second domain entirely with gaps in the other sequence. The alignments are also visualized in third row of Figure 11

**Figure 13:**
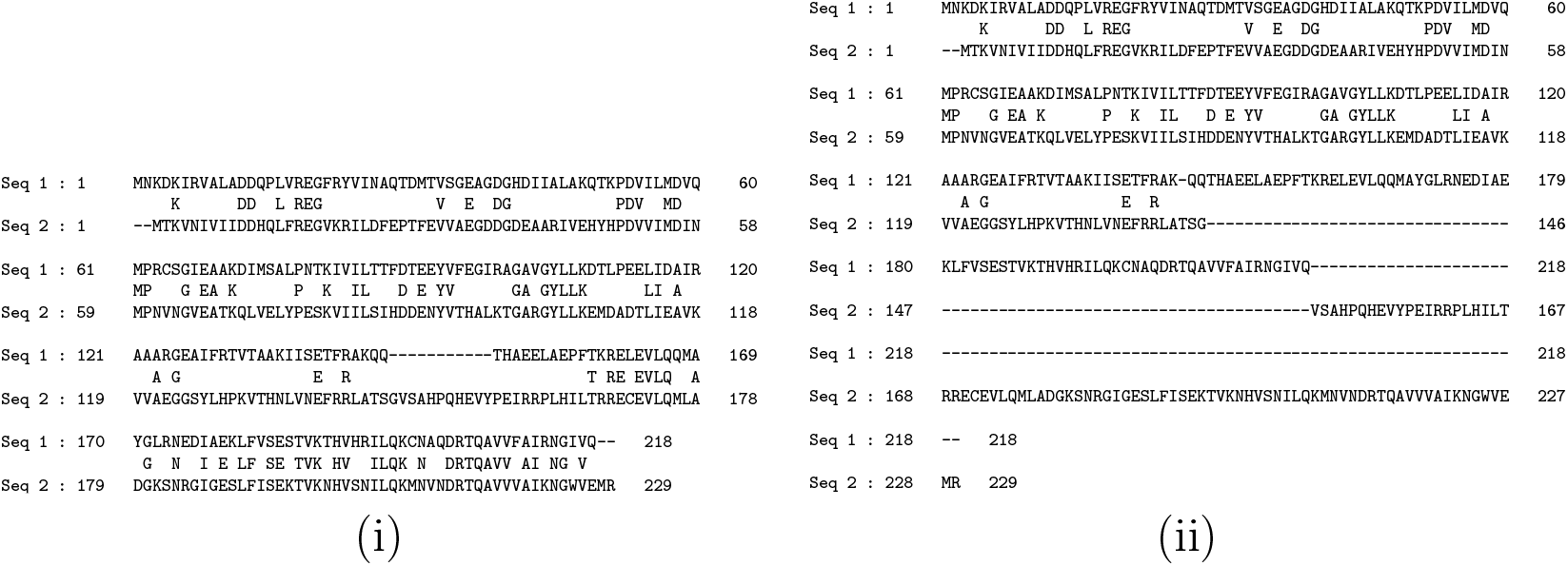
The sequence alignments for YxjL (`Seq 1`) and DegU (`Seq 2`): (i) reference and Ankh score and (ii) AF3US. (Only two alignments are shown because the Ankh-score alignment is identical with the reference.)

## 5. Experimentally determined structures

One aspect that needs to be investigated is that of experimentally determined structures. They are expected to be superior to the AlphaFold-predicted structures, implying that the sequence alignments produced by US-aligning experimentally determined structures are expected to be better as well. We haver attempted to clarify this aspect, with limited success. The problem is the lack of experimentally determined structures for the exact sequences in our MSAs. The sequences must be identical, otherwise the distances cannot be computed. Through considerable manual effort, we identified and tested a few protein pairs with suitable experimentally determined structures. We identified 11 suitable sequences from the cd14765 domain, giving us, with the four distances, 220 tests. The results are summarized in Table 2. The Ankh-score method is still clearly superior to both AF3US and experimental structures. The unexpected result is AF3US against experimental (aligned with US-align). AF3US wins 41.82% of tests, whereas the experimental structures, US-aligned, win only 35.45% of tests, with the remaining 22.73% being ties. Any conclusion drawn from this comparison is not reliable, due to the small number of samples, the fact that they all came from the same domain, and the fact that not even reference MSAs are perfect. Yet, the results presented are intriguing and sufficient to justify further investigation.

**Table 2:** Comparing US-aligned experimentally determined structures with Ankh-score and AF3US. A total of 220 tests from the cd14765 domain were used.

|  |  |  |  |
| --- | --- | --- | --- |
| AF3US vs Experimental | AF3US wins<br>92 | Experimental wins<br>78 | equal<br>50 |
| Ankh-score vs Experimental | Ankh-score wins<br>150 | Experimental wins<br>30 | equal<br>40 |
| Ankh-score vs AF3US | Ankh-score wins<br>150 | AF3US wins<br>27 | equal<br>43 |

## 6. Conclusion

Protein sequence alignments are fundamental in bioinformatics and biology because they provide explicit residue-to-residue correspondence, essential for tracing evolutionary relationships, detecting conserved motifs, and building profile databases. Alignments uniquely reveal positional conservation across homologs, grounding molecular biology in evolutionary context. Their function is essential and improvements to the quality of computational alignments are very important. We investigated here two powerful candidates for improving protein sequence alignments: AlphaFold and protein language models.

AlphaFold marked a breakthrough in biology, providing a much better solution to the decades-long protein structure prediction challenge. By providing near-experimental accuracy at scale, it transforms structural biology, accelerating drug discovery, enzyme design, and functional annotation. Alignment of its predicted structures is considered as a challenger for traditional alignments.

The appearance of protein language models introduced a transformative shift in bioinformatics, enabling rich vector representations of sequences learned from massive unlabelled datasets. Their embeddings capture subtle biochemical and structural patterns, allowing functional prediction, remote homology detection, and clustering beyond sequence identity. The cosine similarity of embeddings has been shown to produce improved sequence alignments.

We compared the two proposed improvements thoroughly and the result is the clear domination of the Ankh-score as the current best method for computing protein sequence alignments. This result has the important implication that protein language models, especially Ankh, may possess certain information that is not available in the AlphaFold3-predicted structures, which requires further investigation.

We considered very briefly experimental structures, which are expected to surpass AF3US. Very limited testing indicates, rather unexpectedly, the opposite. This issue clearly deserves more investigation.

## Acknowledgements

All our computations were performed on Digital Research Alliance of Canada servers.

## Availability

The alignment software is freely available as a web server at e-score. csd.uwo.ca and as source code at github.com/lucian-ilie/E-score.

## Competing interests

No competing interest is declared.

## Author contributions

J.M. selected the data, performed all tests, including installing and running the necessary libraries, and contributed to the methodology and analysis. K.R. performed the detailed biological analysis of case studies. G.B.G. advised on choosing the data and contributed to the biological analysis. L.I. proposed the study, designed the methodology, analyzed the results, supervised the work and wrote the manuscript.

## Funding

This work was supported by NSERC Discovery RGPIN-2021-03978 to and RGPIN-2020-05733 to G.B.G.

## Notes

### Competing Interest Statement

The authors have declared no competing interest.

### Summary of Updates

Changes were implemented: - the software is available as web server and source code at GitHub - gap penalty sensitivity analysis provided - Ankh has been proven the best protein language model for computing alignments by extensive comparison with ProtT5, ProstT5, and ESM-C - Ankh has been proven superior to other PLM-based methods such as PEbA and pLM-BLAST - US-align has been proven the best structural aligner for our task by thorough comparison with DALI and Foldseek - while Ankh is clearly the top choice, the other top PLMs considered have bee proven superior as well to the US-align + AlphaFold3 method, indicating that PLMs in general may possess certain information that is not available in the AlphaFold3-predicted structures.

