## Supplementary Tables and Figures for "Ankh-score produces better sequence alignments than AlphaFold3"

– Supplementary material –

Julia Malec, Karina Rusen, G. Brian Golding, Lucian Ilie*

April 2, 2026

#### 1 Datasets

Supplementary Table 1: BAliBASE domains.

| Domain | Sequences | Tests | Avg. length | Avg. identity (%) | Avg. min TM-score | Avg. max TM-score |
|---|---|---|---|---|---|---|
| BB20008 | 1,431 | 1,023,165 | 522.96 | 12.71 | 0.30 | 0.55 |
| BB20015 | 630 | 198,135 | 193.61 | 21.33 | 0.35 | 0.48 |
| BB30014 | 946 | 446,985 | 217.32 | 23.93 | 0.78 | 0.82 |
| BB30016 | 666 | 221,445 | 291.30 | 11.75 | 0.23 | 0.40 |
| BB40017 | 861 | 370,230 | 224.05 | 16.73 | 0.43 | 0.70 |
| BB40036 | 406 | 82,215 | 338.45 | 33.33 | 0.75 | 0.91 |
| BB50013 | 153 | 11,628 | 283.67 | 32.92 | 0.80 | 0.89 |
| BBS20005 | 741 | 274,170 | 341.13 | 38.96 | 0.88 | 0.90 |
| BBS20007 | 253 | 31,878 | 394.04 | 23.63 | 0.84 | 0.87 |
| BBS20012 | 300 | 44,850 | 206.00 | 36.91 | 0.83 | 0.89 |
| BBS20016 | 351 | 61,425 | 106.30 | 33.71 | 0.86 | 0.90 |
| BBS20017 | 990 | 489,555 | 560.13 | 48.88 | 0.94 | 0.95 |
| BBS20028 | 1,378 | 948,753 | 252.30 | 46.81 | 0.89 | 0.95 |
| BBS20029 | 406 | 82,215 | 58.55 | 36.59 | 0.73 | 0.74 |
| BBS20032 | 1,081 | 583,740 | 100.17 | 40.28 | 0.86 | 0.90 |
| BBS20038 | 741 | 274,170 | 98.92 | 43.42 | 0.87 | 0.92 |
| BBS30002 | 465 | 107,880 | 471.74 | 23.13 | 0.73 | 0.87 |
| BBS30014 | 946 | 446,985 | 203.36 | 25.60 | 0.82 | 0.84 |
| BOSX049 | 153 | 11,628 | 97.61 | 27.85 | 0.52 | 0.55 |
| BOSX122 | 153 | 11,628 | 530.33 | 52.62 | 0.93 | 0.94 |
| Total | 13,051 | 5,722,680 | 274.60 | 31.55 | 0.72 | 0.80 |

Supplementary Table 2: CDD domains.

| Domain | Sequences | Tests | Avg. length | Avg. identity (%) | Avg. min TM-score | Avg. max TM-score |
|---|---|---|---|---|---|---|
| cd00001 | 300 | 44,850 | 180.40 | 25.43 | 0.79 | 0.91 |
| cd00005 | 406 | 82,215 | 1112.62 | 9.04 | 0.35 | 0.53 |
| cd00025 | 105 | 5,460 | 477.87 | 14.23 | 0.75 | 0.78 |
| cd00103 | 325 | 52,650 | 373.88 | 10.47 | 0.35 | 0.50 |
| cd00148 | 325 | 52,650 | 140.50 | 25.63 | 0.77 | 0.86 |
| cd00173 | 595 | 176,715 | 82.23 | 14.59 | 0.58 | 0.64 |
| cd00214 | 276 | 37,950 | 780.83 | 12.70 | 0.61 | 0.71 |
| cd00454 | 1,485 | 1,101,870 | 795.65 | 5.22 | 0.27 | 0.60 |
| cd01040 | 1,326 | 878,475 | 352.73 | 11.87 | 0.52 | 0.80 |
| cd05303 | 666 | 221,445 | 312.19 | 37.25 | 0.90 | 0.93 |
| cd06444 | 325 | 52,650 | 707.50 | 19.55 | 0.66 | 0.80 |
| cd06609 | 210 | 21,945 | 528.90 | 25.91 | 0.50 | 0.70 |
| cd12124 | 210 | 21,945 | 195.81 | 50.00 | 0.96 | 0.97 |
| cd12176 | 465 | 107,880 | 435.84 | 38.31 | 0.85 | 0.95 |
| cd12186 | 1,081 | 583,740 | 332.00 | 37.00 | 0.94 | 0.95 |
| cd14765 | 78 | 3,003 | 141.92 | 41.07 | 0.91 | 0.93 |
| cd15492 | 903 | 407,253 | 46.70 | 46.12 | 0.89 | 0.91 |
| cd19754 | 105 | 5,460 | 397.73 | 28.09 | 0.91 | 0.95 |
| cd19756 | 595 | 176,715 | 39.00 | 27.01 | 0.69 | 0.71 |
| cd21762 | 595 | 176,715 | 529.86 | 4.04 | 0.11 | 0.30 |
| Total | 10,376 | 4,211,586 | 398.21 | 24.18 | 0.67 | 0.77 |

### 2 Gap penalty sensitivity analysis

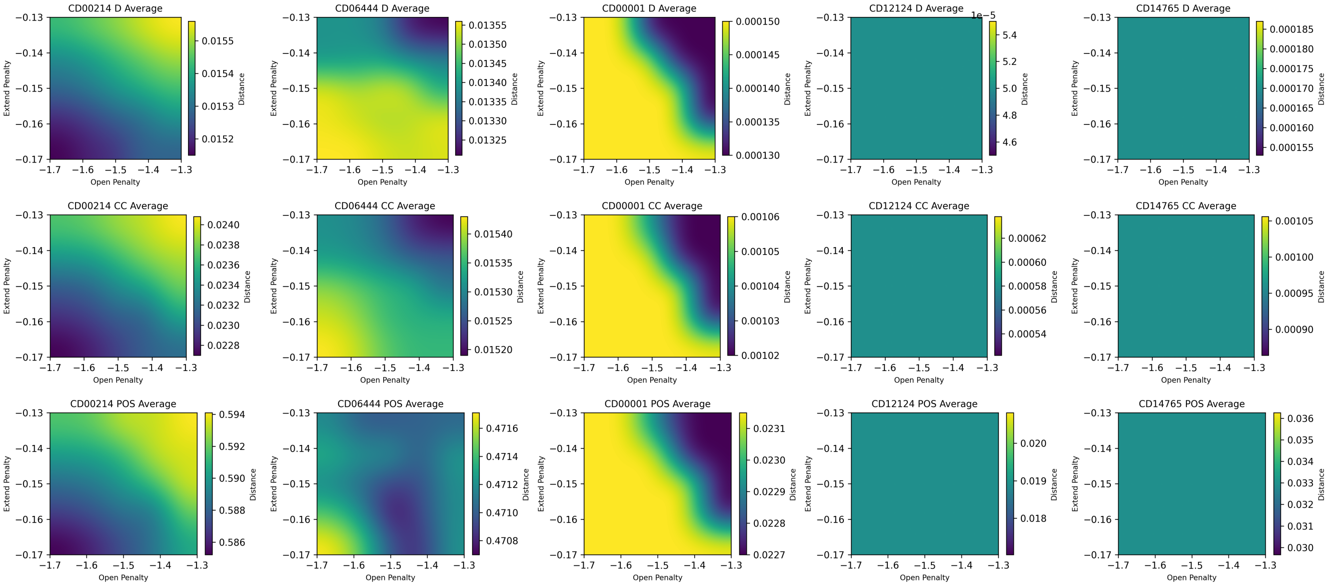


Supplementary Figure 1: Heat maps showing the variation of the $d_d$, $d_{cc}$, and $d_{pos}$ distances between the Ankh-score alignments and the reference for several domains.

### 3 Results for the $\mathrm{d_d}$, $\mathrm{d_{cc}}$, and $\mathrm{d_{pos}}$ distances

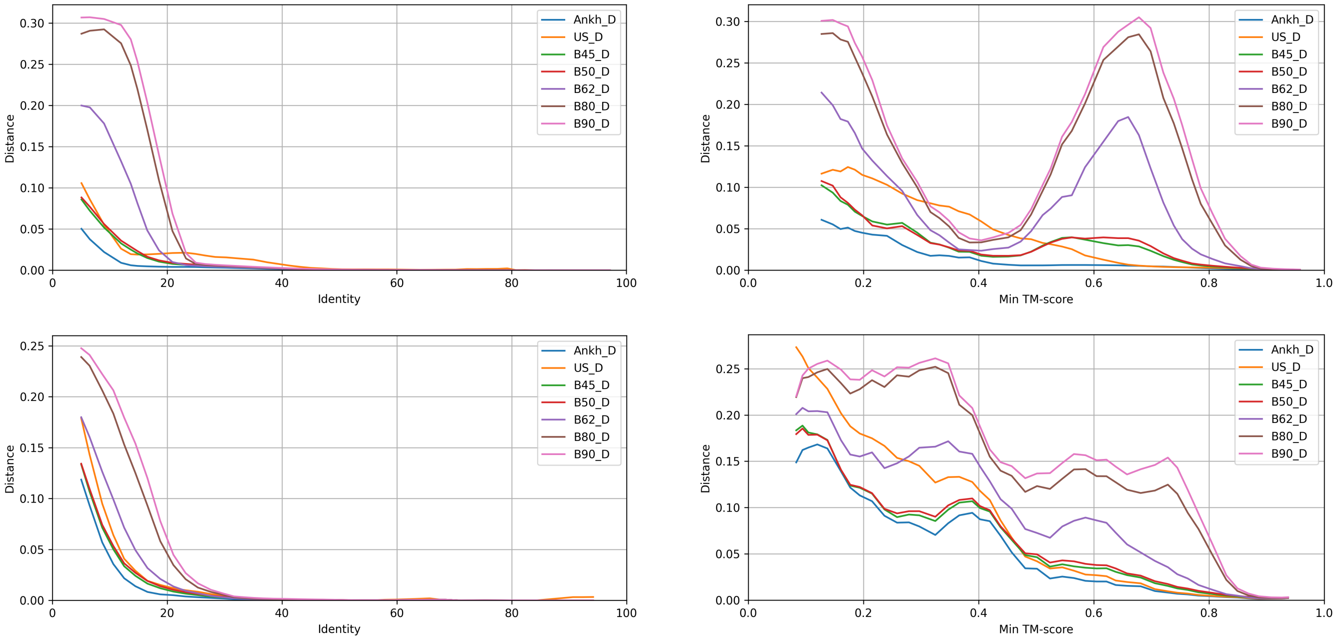


Supplementary Figure 2: All methods compared on all tests for the $\mathrm{d_d}$ distance. The plots in the first row are for BAliBASE datasets, while those is the second row are for CDD. In the first column, the scores are sorted by the identity level of the sequences, while in the second column, they are sorted by the minimum TM-score.

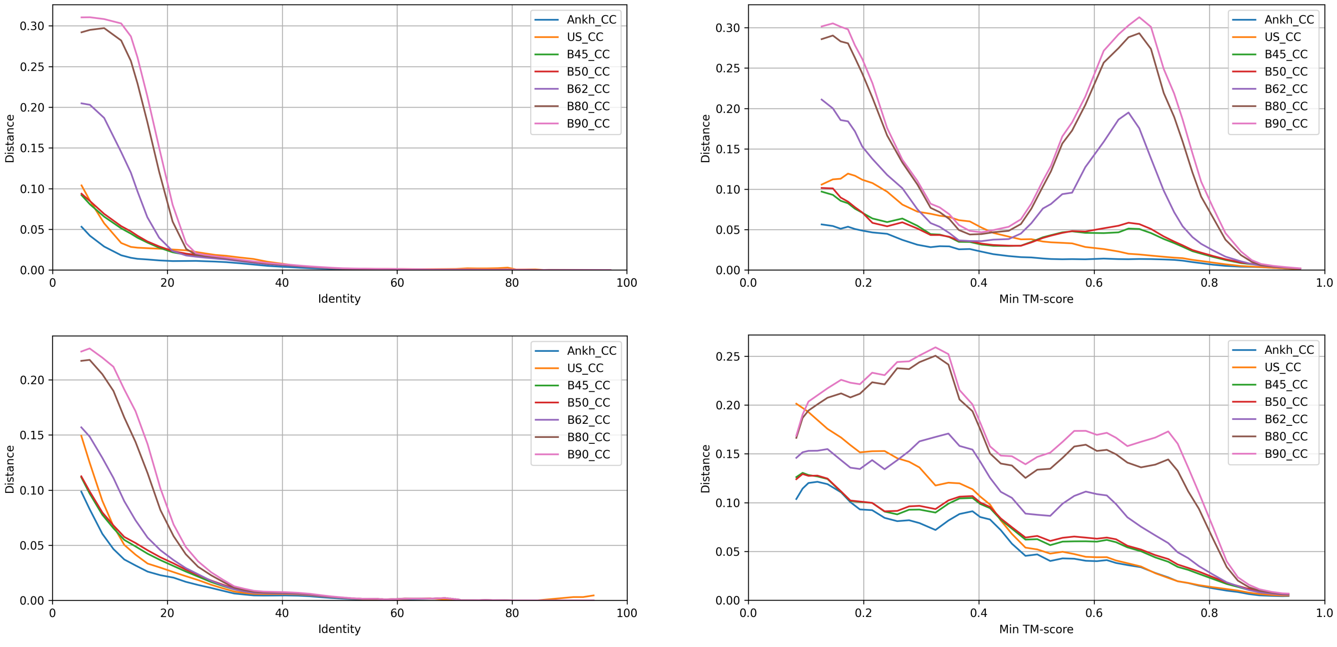


Supplementary Figure 3: All methods compared on all tests for the $\mathrm{d_{cc}}$ distance. The plots in the first row are for BAliBASE datasets, while those is the second row are for CDD. In the first column, the scores are sorted by the identity level of the sequences, while in the second column, they are sorted by the minimum TM-score.

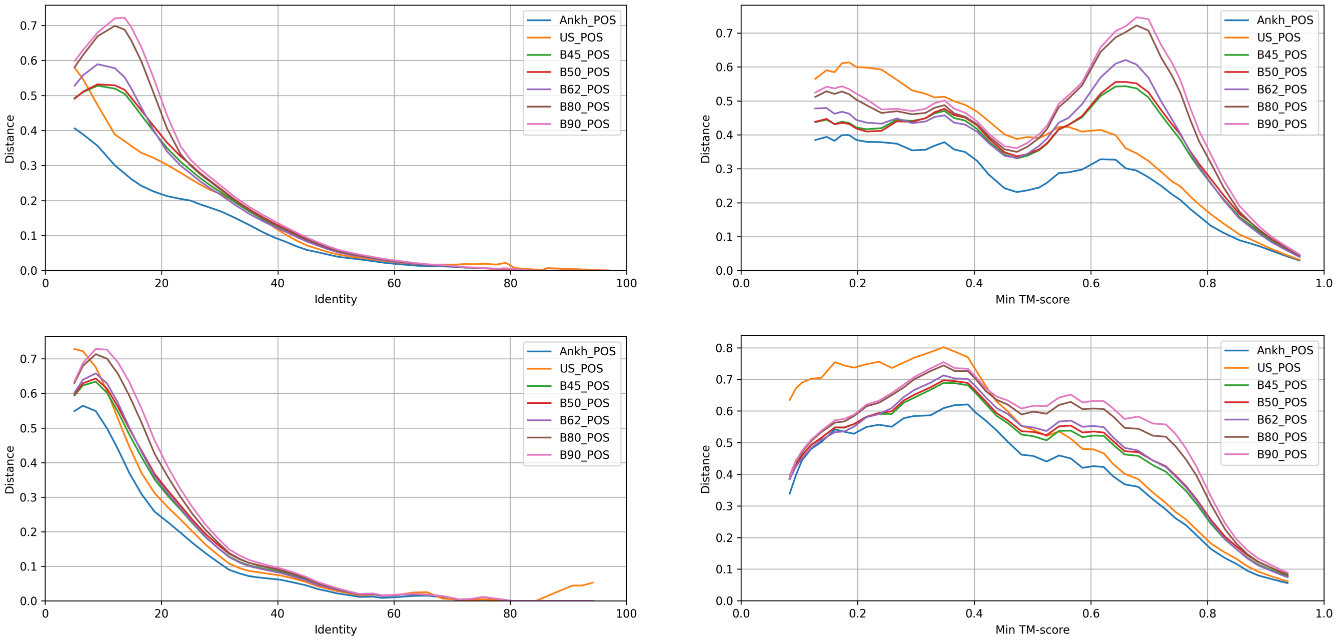


Supplementary Figure 4: All methods compared on all tests for the $d_{pos}$ distance. The plots in the first row are for BAliBASE datasets, while those is the second row are for CDD. In the first column, the scores are sorted by the identity level of the sequences, while in the second column, they are sorted by the minimum TM-score.

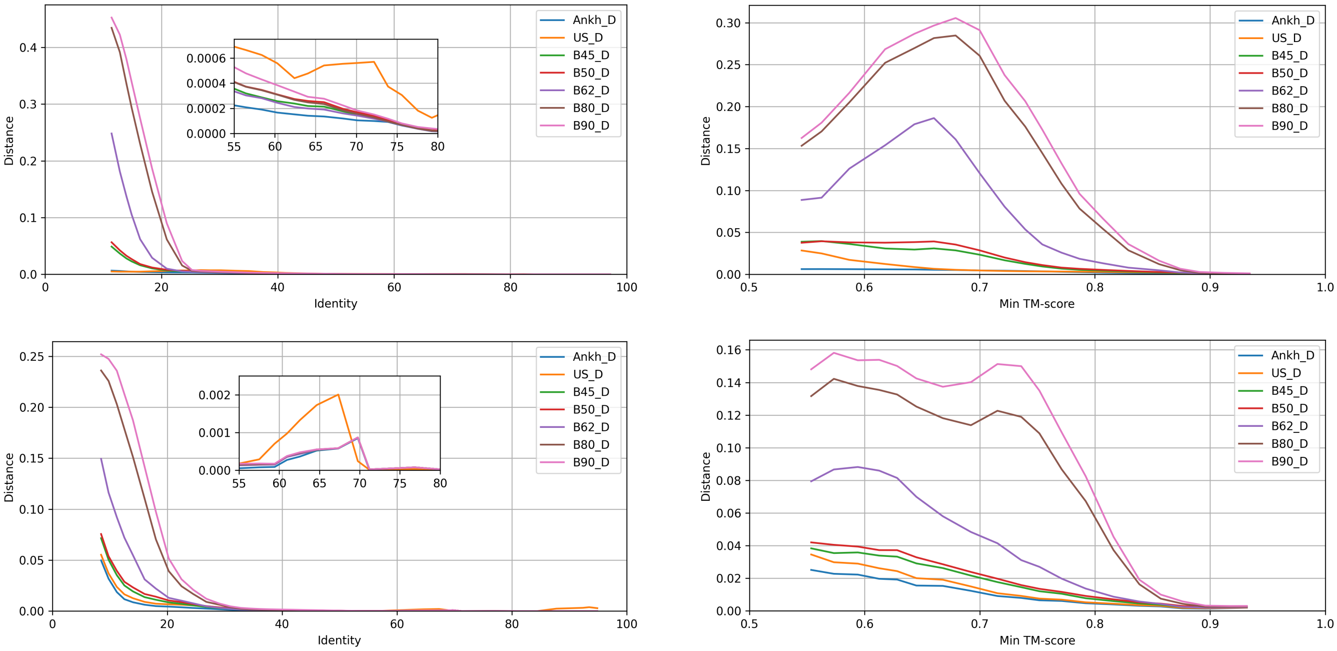


Supplementary Figure 5: All methods are compared on tests with minimum TM-score > 0.5 for the $d_d$ distance. The plots in the first row are for BAliBASE datasets, while those is the second row are for CDD. In the first column, the scores are sorted by the identity level of the sequences, while in the second column, they are sorted by the minimum TM-score.

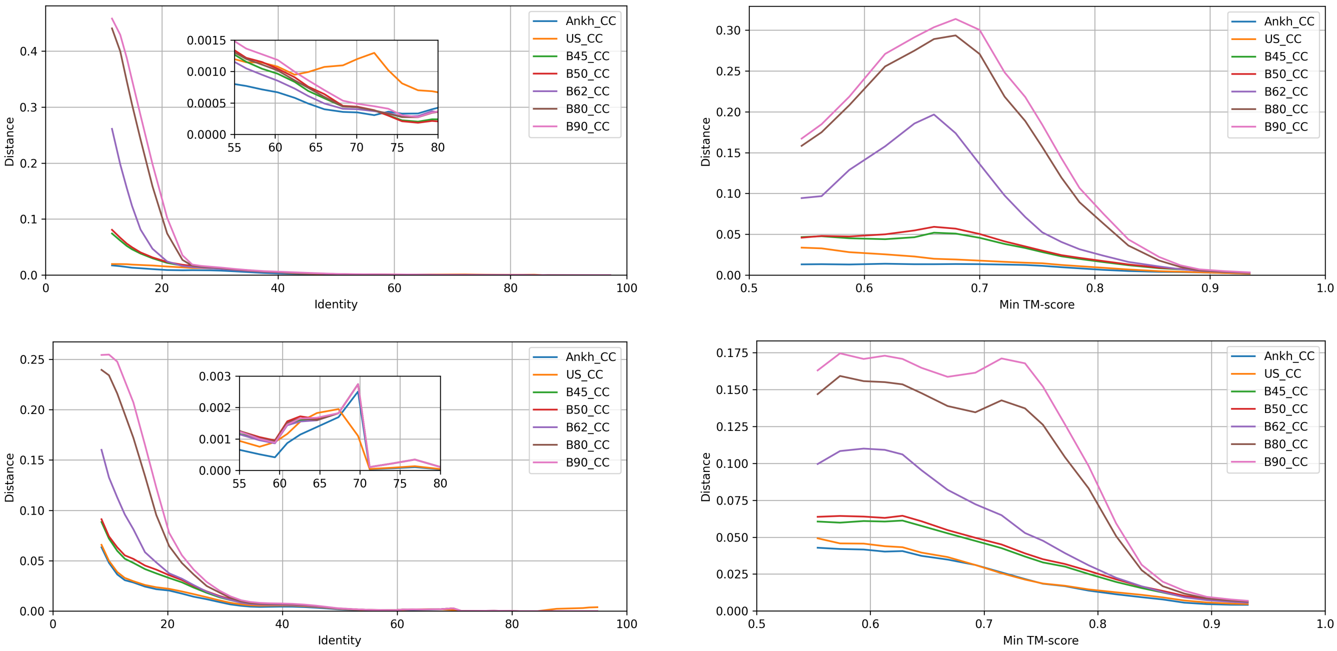


Supplementary Figure 6: All methods are compared on tests with minimum TM-score > 0.5 for the $d_{cc}$ distance. The plots in the first row are for BAliBASE datasets, while those is the second row are for CDD. In the first column, the scores are sorted by the identity level of the sequences, while in the second column, they are sorted by the minimum TM-score.

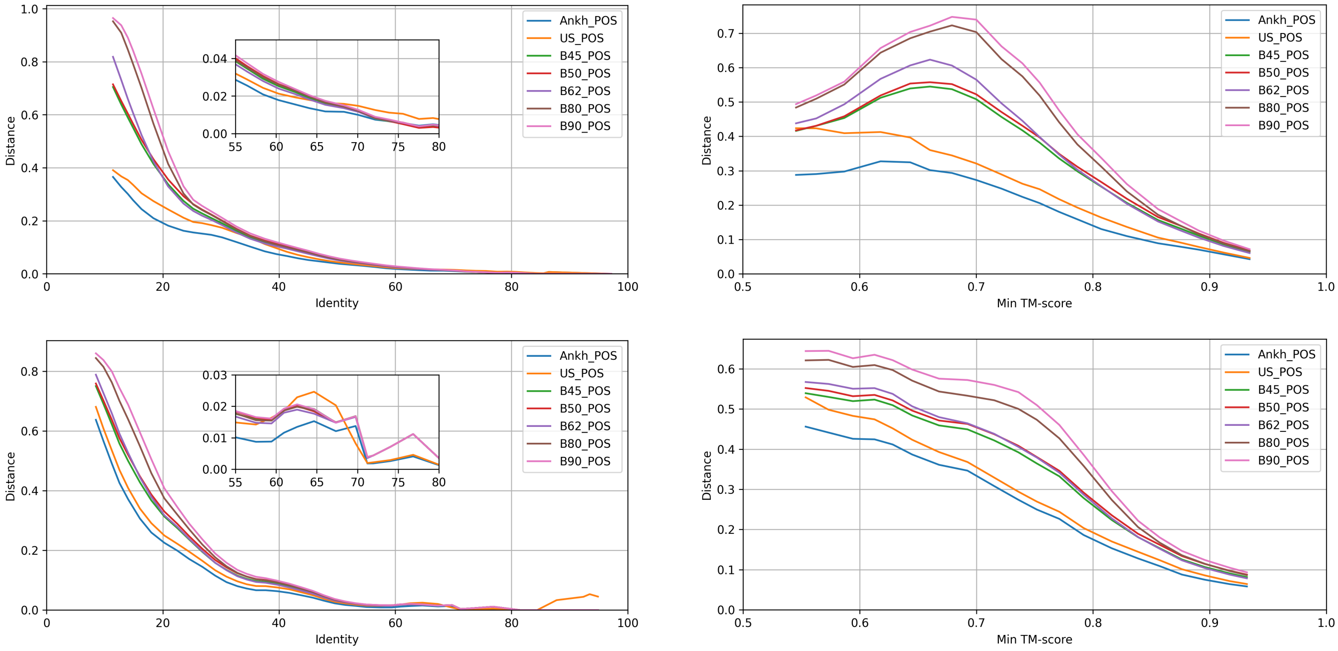


Supplementary Figure 7: All methods are compared on tests with minimum TM-score > 0.5 for the $d_{pos}$ distance. The plots in the first row are for BAliBASE datasets, while those is the second row are for CDD. In the first column, the scores are sorted by the identity level of the sequences, while in the second column, they are sorted by the minimum TM-score.

### 4 Head-to-head comparison – summary of results

Supplementary Table 3: Head-to-head comparison – summary of results for all tests. For each of the three method pairs compared, we give the total number of domains won and the total number and percentage of tests won, for each database, BAliBASE and CDD, as well as combined. The results are given separately for each of the four distances considered. "null" gives the number of domains with p-values higher than 0.01 and "equal" gives the fraction of tests for which the performances are the same.

| Dataset | $d_{ia}$ | | | $d_d$ | | | $d_{cc}$ | | | $d_{pos}$ | | |
|---|---|---|---|---|---|---|---|---|---|---|---|---|
| **Ankh vs AF3US** | | | | | | | | | | | | |
| Domains won | Ankh | AF3US | null | Ankh | AF3US | null | Ankh | AF3US | null | Ankh | AF3US | null |
| BAliBASE | **14** | 3 | 3 | **13** | 3 | 4 | **15** | 3 | 2 | **16** | 3 | 1 |
| CDD | **17** | 2 | 1 | **17** | 1 | 2 | **16** | 1 | 3 | **18** | 1 | 1 |
| Total | **31** | 5 | 4 | **30** | 4 | 6 | **31** | 4 | 5 | **34** | 4 | 2 |
| Tests won | Ankh | AF3US | equal | Ankh | AF3US | equal | Ankh | AF3US | equal | Ankh | AF3US | equal |
| BAliBASE | **6897** | 2792 | 799 | **6695** | 3289 | 1250 | **7745** | 2893 | 1079 | **7811** | 2805 | 1354 |
| CDD | **6073** | 2545 | 1163 | **6169** | 1909 | 1037 | **6295** | 2109 | 947 | **6435** | 1921 | 1425 |
| Total | **12970** | 5337 | 1962 | **12864** | 5198 | 2287 | **14040** | 5002 | 2026 | **14246** | 4726 | 2779 |
| Tests won (%) | Ankh | AF3US | equal | Ankh | AF3US | equal | Ankh | AF3US | equal | Ankh | AF3US | equal |
| BAliBASE | **66.05** | 26.80 | 7.15 | **60.52** | 29.24 | 10.24 | **65.38** | 25.94 | 8.68 | **65.04** | 23.91 | 11.05 |
| CDD | **67.16** | 22.29 | 10.55 | **74.02** | 16.11 | 9.87 | **71.01** | 19.86 | 9.13 | **70.22** | 17.40 | 12.38 |
| Total | **66.61** | 24.55 | 8.85 | **67.27** | 22.67 | 10.05 | **68.19** | 22.90 | 8.91 | **67.63** | 20.66 | 11.72 |
| **AF3US vs Blosum45** | | | | | | | | | | | | |
| Domains won | AF3US | Blosum45 | null | AF3US | Blosum45 | null | AF3US | Blosum45 | null | AF3US | Blosum45 | null |
| BAliBASE | **14** | 4 | 2 | **11** | 6 | 3 | **13** | 5 | 2 | **16** | 4 | 0 |
| CDD | **11** | 8 | 1 | **8** | 7 | 5 | **11** | 7 | 2 | **11** | 7 | 2 |
| Total | **25** | 12 | 3 | **19** | 13 | 8 | **24** | 12 | 4 | **27** | 11 | 2 |
| Tests won | AF3US | Blosum45 | equal | AF3US | Blosum45 | equal | AF3US | Blosum45 | equal | AF3US | Blosum45 | equal |
| BAliBASE | **7033** | 3946 | 1000 | **5872** | 4841 | 860 | **7094** | 4073 | 812 | **7723** | 4134 | 1194 |
| CDD | **4963** | 4089 | 421 | **4192** | 4001 | 307 | **4597** | 4135 | 276 | **4394** | 3928 | 686 |
| Total | **11996** | 8035 | 1421 | **10064** | 8842 | 1167 | **11691** | 8208 | 1088 | **12117** | 8062 | 1880 |
| Tests won (%) | AF3US | Blosum45 | equal | AF3US | Blosum45 | equal | AF3US | Blosum45 | equal | AF3US | Blosum45 | equal |
| BAliBASE | **59.20** | 32.97 | 7.83 | **53.42** | 40.07 | 6.51 | **60.28** | 33.89 | 5.83 | **60.10** | 31.34 | 8.56 |
| CDD | **49.00** | 45.33 | 5.67 | 45.51 | **50.17** | 4.31 | **49.42** | 46.34 | 4.24 | **48.97** | 43.65 | 7.38 |
| Total | **54.10** | 39.15 | 6.75 | **49.47** | 45.12 | 5.41 | **54.85** | 40.11 | 5.03 | **54.54** | 37.49 | 7.97 |
| **Ankh-score vs Blosum45** | | | | | | | | | | | | |
| Domains won | Ankh | Blosum45 | null | Ankh | Blosum45 | null | Ankh | Blosum45 | null | Ankh | Blosum45 | null |
| BAliBASE | **17** | 1 | 2 | **16** | 1 | 3 | **17** | 1 | 2 | **18** | 0 | 2 |
| CDD | **15** | 2 | 3 | **20** | 0 | 0 | **18** | 1 | 1 | **19** | 0 | 1 |
| Total | **32** | 3 | 5 | **36** | 1 | 3 | **35** | 2 | 3 | **37** | 0 | 3 |
| Tests won | Ankh | Blosum45 | equal | Ankh | Blosum45 | equal | Ankh | Blosum45 | equal | Ankh | Blosum45 | equal |
| BAliBASE | **9433** | 1732 | 1433 | **8664** | 2055 | 1138 | **9344** | 1734 | 1079 | **9468** | 1362 | 1327 |
| CDD | **5920** | 1487 | 1169 | **7158** | 1932 | 1286 | **6943** | 1981 | 1242 | **6330** | 1149 | 1412 |
| Total | **15353** | 3219 | 2602 | **15822** | 3987 | 2424 | **16287** | 3715 | 2321 | **15798** | 2511 | 2739 |
| Tests won (%) | Ankh | Blosum45 | equal | Ankh | Blosum45 | equal | Ankh | Blosum45 | equal | Ankh | Blosum45 | equal |
| BAliBASE | **75.60** | 12.86 | 11.54 | **72.74** | 17.39 | 9.87 | **76.89** | 13.89 | 9.22 | **77.91** | 10.87 | 11.23 |
| CDD | **70.17** | 17.63 | 12.20 | **72.79** | 16.29 | 10.92 | **70.03** | 19.06 | 10.92 | **73.98** | 12.93 | 13.09 |
| Total | **72.88** | 15.25 | 11.87 | **72.77** | 16.84 | 10.39 | **73.46** | 16.47 | 10.07 | **75.95** | 11.90 | 12.16 |

Supplementary Table 4: Head-to-head comparison – summary of results for tests with minimum TM-score > 0.5. For each of the three method pairs compared, we give the total number of domains won and the total number and percentage of tests won, for each database, BAliBASE and CDD, as well as combined. The results are given separately for each of the four distances considered. "null" gives the number of domains with p-values higher than 0.01 and "equal" gives the fraction of tests for which the performances are the same.

| Dataset | $d_{ia}$ | | | $d_d$ | | | $d_{cc}$ | | | $d_{pos}$ | | |
|---|---|---|---|---|---|---|---|---|---|---|---|---|
| Ankh-score vs AF3US | | | | | | | | | | | | |
| Domains won | Ankh | AF3US | null | Ankh | AF3US | null | Ankh | AF3US | null | Ankh | AF3US | null |
| BAliBASE | **14** | 2 | 4 | **11** | 4 | 5 | **14** | 3 | 3 | **15** | 3 | 2 |
| CDD | **16** | 2 | 2 | **15** | 1 | 4 | **13** | 0 | 7 | **16** | 1 | 3 |
| Total | **30** | 4 | 6 | **26** | 5 | 9 | **27** | 3 | 10 | **31** | 4 | 5 |
| Tests won | Ankh | AF3US | equal | Ankh | AF3US | equal | Ankh | AF3US | equal | Ankh | AF3US | equal |
| BAliBASE | **4227** | 2315 | 756 | **3966** | 2859 | 1211 | **5081** | 2390 | 1048 | **5217** | 2283 | 1272 |
| CDD | **3539** | 1583 | 1115 | **3336** | 1061 | 940 | **3214** | 1054 | 853 | **3442** | 1212 | 1132 |
| Total | **7766** | 3898 | 1871 | **7302** | 3920 | 2151 | **8295** | 3444 | 1901 | **8659** | 3495 | 2404 |
| Tests won (%) | Ankh | AF3US | equal | Ankh | AF3US | equal | Ankh | AF3US | equal | Ankh | AF3US | equal |
| BAliBASE | **65.54** | 26.31 | 8.16 | **59.04** | 29.41 | 11.55 | **65.60** | 24.63 | 9.77 | **65.23** | 22.53 | 12.24 |
| CDD | **65.20** | 23.18 | 11.62 | **74.17** | 15.63 | 10.20 | **69.04** | 20.39 | 10.58 | **67.92** | 17.77 | 14.31 |
| Total | **65.37** | 24.74 | 9.89 | **66.60** | 22.52 | 10.87 | **67.32** | 22.51 | 10.17 | **66.57** | 20.15 | 13.28 |
| AF3US vs Blosum45 | | | | | | | | | | | | |
| Domains won | AF3US | Blosum45 | null | AF3US | Blosum45 | null | AF3US | Blosum45 | null | AF3US | Blosum45 | null |
| BAliBASE | **12** | 6 | 2 | **11** | 6 | 3 | **13** | 5 | 2 | **14** | 3 | 3 |
| CDD | **10** | 6 | 4 | **10** | 4 | 6 | **11** | 4 | 5 | **11** | 4 | 5 |
| Total | **22** | 12 | 6 | **21** | 10 | 9 | **24** | 9 | 7 | **25** | 7 | 8 |
| Tests won | AF3US | Blosum45 | equal | AF3US | Blosum45 | equal | AF3US | Blosum45 | equal | AF3US | Blosum45 | equal |
| BAliBASE | **6492** | 2113 | 899 | **5434** | 2865 | 847 | **6408** | 2291 | 805 | **6424** | 1849 | 985 |
| CDD | **3543** | 1541 | 375 | **3179** | 1609 | 279 | **3348** | 1631 | 245 | **3394** | 1210 | 408 |
| Total | **10035** | 3654 | 1274 | **8613** | 4474 | 1126 | **9756** | 3922 | 1050 | **9818** | 3059 | 1393 |
| Tests won (%) | AF3US | Blosum45 | equal | AF3US | Blosum45 | equal | AF3US | Blosum45 | equal | AF3US | Blosum45 | equal |
| BAliBASE | **60.00** | 32.99 | 7.01 | **53.39** | 39.77 | 6.84 | **59.85** | 33.97 | 6.18 | **62.36** | 29.63 | 8.01 |
| CDD | **53.36** | 39.87 | 6.77 | **48.45** | 46.58 | 4.97 | **56.29** | 38.62 | 5.09 | **55.96** | 35.74 | 8.30 |
| Total | **56.68** | 36.43 | 6.89 | **50.92** | 43.17 | 5.91 | **58.07** | 36.30 | 5.63 | **59.16** | 32.69 | 8.16 |
| Ankh-score vs Blosum45 | | | | | | | | | | | | |
| Domains won | Ankh | Blosum45 | null | Ankh | Blosum45 | null | Ankh | Blosum45 | null | Ankh | Blosum45 | null |
| BAliBASE | **18** | 0 | 2 | **18** | 0 | 2 | **17** | 0 | 3 | **18** | 0 | 2 |
| CDD | **15** | 0 | 5 | **17** | 1 | 2 | **15** | 1 | 4 | **18** | 0 | 2 |
| Total | **33** | 0 | 7 | **35** | 1 | 4 | **32** | 1 | 7 | **36** | 0 | 4 |
| Tests won | Ankh | Blosum45 | equal | Ankh | Blosum45 | equal | Ankh | Blosum45 | equal | Ankh | Blosum45 | equal |
| BAliBASE | **7080** | 1013 | 1377 | **6614** | 1512 | 1344 | **6928** | 1056 | 1042 | **7048** | 791 | 1198 |
| CDD | **4535** | 814 | 1144 | **4761** | 812 | 1119 | **4602** | 925 | 1071 | **4828** | 661 | 1238 |
| Total | **11615** | 1827 | 2521 | **11375** | 2324 | 2463 | **11530** | 1981 | 2113 | **11876** | 1452 | 2436 |
| Tests won (%) | Ankh | Blosum45 | equal | Ankh | Blosum45 | equal | Ankh | Blosum45 | equal | Ankh | Blosum45 | equal |
| BAliBASE | **75.81** | 11.48 | 12.71 | **70.69** | 16.90 | 12.40 | **76.85** | 12.08 | 11.07 | **78.94** | 9.10 | 11.96 |
| CDD | **70.56** | 13.96 | 15.48 | **73.89** | 12.67 | 13.44 | **69.86** | 16.21 | 13.93 | **73.29** | 10.93 | 15.78 |
| Total | **73.18** | 12.72 | 14.10 | **72.29** | 14.79 | 12.92 | **73.36** | 14.14 | 12.50 | **76.11** | 10.01 | 13.87 |
